# Expanding the molecular grammar of polar residues and arginine in FUS prion-like domain phase separation and aggregation

**DOI:** 10.1101/2024.02.15.580391

**Authors:** Noah Wake, Shuo-Lin Weng, Tongyin Zheng, Szu-Huan Wang, Valentin Kirilenko, Jeetain Mittal, Nicolas L Fawzi

**Affiliations:** Therapeutic Sciences Graduate Program, Brown University, Providence, RI 02912; Department of Chemistry, Texas A&M University, College Station, TX 77843; Department of Molecular Biology, Cell Biology & Biochemistry, Brown University, Providence, RI 02912; Artie McFerrin Department of Chemical Engineering, Texas A&M University, College Station, TX 77843; Interdisciplinary Graduate Program in Genetics and Genomics, Texas A&M University, College Station, TX 77843

**Author notes:** NW and SLW contributed equally.

## Abstract

A molecular grammar governing low-complexity prion-like domains phase separation (PS) has been proposed based on mutagenesis experiments that identified tyrosine and arginine as primary drivers of phase separation via aromatic-aromatic and aromatic-arginine interactions. Here we show that additional residues make direct favorable contacts that contribute to phase separation, highlighting the need to account for these contributions in PS theories and models. We find that tyrosine and arginine make important contacts beyond only tyrosine-tyrosine and tyrosine-arginine, including arginine-arginine contacts. Among polar residues, glutamine in particular contributes to phase separation with sequence/position-specificity, making contacts with both tyrosine and arginine as well as other residues, both before phase separation and in condensed phases. For glycine, its flexibility, not its small solvation volume, favors phase separation by allowing favorable contacts between other residues and inhibits the liquid-to-solid (LST) transition. Polar residue types also make sequence-specific contributions to aggregation that go beyond simple rules, which for serine positions is linked to formation of an amyloid-core structure by the FUS low-complexity domain. Hence, here we propose a revised molecular grammar expanding the role of arginine and polar residues in prion-like domain protein phase separation and aggregation.

## Introduction

The spontaneous demixing of biomolecules is stabilized by intermolecular contacts via several interaction modes which contribute to the physical properties of the condensed phase^1–3^. While biomolecular condensates have complex compositions in cells, proteins with domains of low sequence complexity and prion-like residue composition, i.e. resembling the polar residue enriched sequences of yeast prion proteins, are highly represented in condensate formation. These low complexity domains are important for the phase separation of these proteins in biochemical experiments and in cells^4,5^, as well as for function. Therefore, the “molecular grammar” governing phase separation of disordered domains has been the topic of intense study^6,7^.

Models to describe biomolecular phase separation have emerged^6,8–15^, including the “stickers and spacers (SaS)” model based on associative polymers^16,17^ where *stickers* or associative motifs are the major determinants for phase separation, and most applications of this model to biomolecular PS assume that the remaining segments of the chain function as *spacers* that only weakly and indirectly modulate phase separation^6^. This model has been successfully applied to the phase separation of many types of associative biopolymers ranging from chains of globular domains and short linear interaction motifs, RNA and RNA-binding domains, and intrinsically disordered domains^18,19^. Alternatively, molecular models based on a continuum of pairwise amino acid interaction strengths (e.g. HPS framework) have also been applied extensively to characterize the phase separation of disordered and multidomain proteins^20,21^. For describing and predicting the sequence-dependent phase separation based on these complementary approaches, a fundamental question emerges: which amino acids, if any, can be ignored from considerations of thermodynamic driving forces of PS?

In the case of an archetypal protein containing a disordered prion-like domain, Fused in Sarcoma (FUS) phase separation, transcriptional activation, and polymerization all depend on tyrosine residues as shown by tyrosine-to-serine mutagenesis experiments^4,22,23^. Extending these observations, mutagenesis-based amino-acid residue substitution experiments combined with theory and low-resolution computational models then were used to support a SaS model for prion-like domain PS, where aromatic residues, especially tyrosine more prominently than phenylalanine, serve as *stickers*^24–26^. Present in RGG motifs found abundantly in several regions of FUS^27^, arginine has also been identified as an *auxiliary sticker*^25^ that mediates contacts specifically with aromatic residues (via cation-π interactions) that are not found for lysine^28^, which has been suggested to destabilize these interactions via three-body effects^25^. These excellent studies have led some to take the view that the molecular grammar of phase separation can simply be understood in terms of aromatic and arginine residues, specifically for FUS tyrosine-tyrosine and tyrosine-arginine interactions.

Yet there is evidence that these residue types, making up only ∼15% of the sequence, are not the only pair contacts that stabilize protein phase separation^29–35^. In domains like FUS LC, the majority of the sequence (78% is made up of polar/small amino acids that have commonly been labeled spacers: serine 26%, glutamine 23%, glycine 17%, threonine 6%, asparagine 3%, alanine 3%. Models derived from NMR experiments and molecular simulation show that many more favorable contacts are formed involving these additional residue types and emphasize a continuum of contributions across many residue types and interaction modes^32,36–40^. In the related prion-like RNA-binding protein TDP-43, we recently showed that hydrophobic amino acids including methionine do contribute important contacts to phase separation, suggesting that the prion-like domain molecular grammar is broader than aromatic and arginine residues^30^. Therefore, the rules for the contribution of the majority of residues in prion-like domains to phase separation remain incompletely characterized.

Although proposed to not play major roles in driving phase separation, polar residues have been proposed to regulate the liquid-to-solid (LST) transitions of phase separating proteins^6,41^. Many proteins containing prion-like domains are associated with neurodegenerative diseases where they form inclusions^42–47^. In test tubes, these proteins can form liquid-like droplets that can convert to solid-like aggregates over time^4,48,49^. This aggregation may occur at the surface of the condensates on the interface between condensed and dispersed phases^50–52^ where the conformations of the proteins may be altered^26,53^. Several additional rules have been suggested, including that glycine is important for maintaining liquidity while serine and glutamine favor the LST transition^6^. Proline residues have also been noted to be crucial for discouraging solid-like aggregation^49,54^. However, the large number of mutations associated with neurodegeneration and cancers^55^ that involve swapping of polar/small residues that do not fall into these rules suggest that the role of these residues in the LST transition is also incompletely known.

Here we probe the contacts formed in FUS phase separation and the role of polar residue types with the aim of testing and refining the molecular grammar for prion-like domain phase separation and LST transitions. To start, we establish FUS LC-RGG1 as a new model sequence of FUS to better test the role of these residues and then compare results between two different contexts (FUS LC vs. FUS LC-RGG1 due to their distinct sequence compositions). We interrogate the network of interactions formed between each residue type in the FUS LC-RGG1 condensed phase using a combination of NMR and molecular dynamics (MD) simulations, probing for the role of different residue types in both the dispersed and condensed phases with direct structural evidence. Subsequently, we present mutagenic studies to test the role of these polar and small residues in phase separation and LST of FUS disordered domains. Together, these studies support revising the established molecular grammar and give insight into the structural details of both prion-like protein phase separation and aggregation.

## Results

### FUS LC-RGG1 model for phase separation

In order to probe the homotypic and heterotypic domain interactions responsible for phase separation of FUS disordered domains, we needed to move beyond our previous work that focused on the isolated FUS LC or mixtures of LC and RGG domains *in trans*^33^. Hence, we characterized the phase separation of FUS LC-RGG1 (residues 1-284) (Fig. 1a). FUS LC and RGG domains are compositionally distinct – the LC is enriched in polar residues including glutamine, serine, tyrosine, and glycine and depleted in charged residues (only two aspartate residues) and non-tyrosine hydrophobic amino acids, while the RGG1 domain has a region with poly-glycine tracts followed by a region enriched in RGG motifs, negatively charged residues, and hydrophobic residues (methionine and phenylalanine). Compared to the isolated FUS LC, FUS LC-RGG1 phase separates more avidly, with a saturation concentration, *C*_sat_, of ∼10 μM, about 10x lower than FUS LC alone under near physiological salt concentration of 150 mM (Fig. 1b). Unlike FUS LC, FUS LC-RGG1 shows reduced phase separation as ionic strength increases consistent with charge screening^56,57^, suggesting that electrostatic interactions play an important role in driving its phase separation. As FUS LC is nearly devoid of charged residues, these data suggest that homotypic RGG1 interactions (i.e. between RGG1 domains) contribute to phase separation.

**Figure 1.**
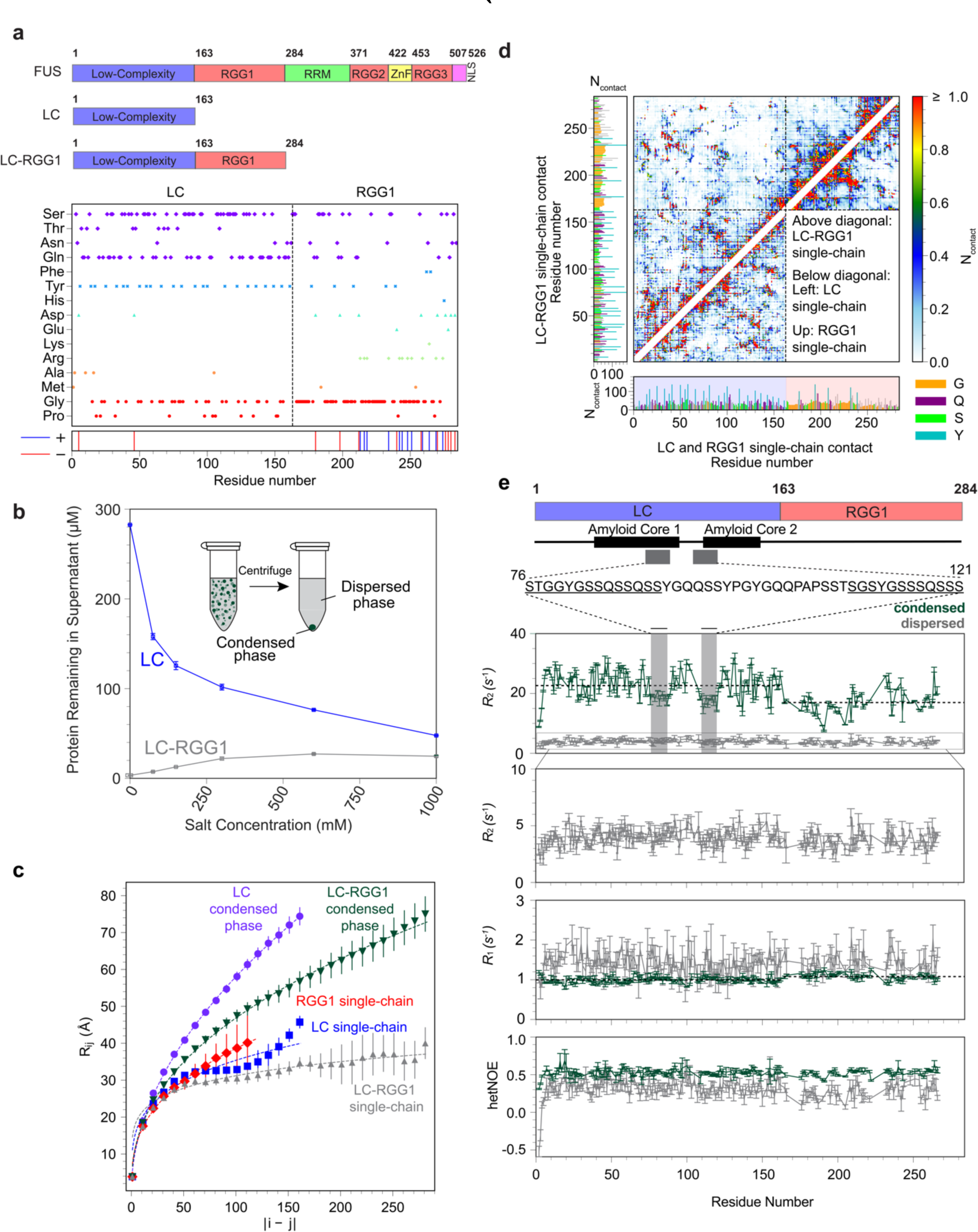
Phase separation of FUS LC-RGG1. **a** FUS domain structure and residue composition of LC-RGG1, showing distinct composition of LC and RGG1 **b** Salt dependence of phase separation (i.e. the saturation concentration) compared for FUS LC and FUS LC-RGG1, measured by the amount of protein remaining in the supernatant. Insert: schematic of the centrifugation experiment to measure the saturation concentration. **c** The average intramolecular distance R_ij_ between the ith and jth residues, calculated from atomistic simulations of isolated monomers (“single chains”) of FUS wild type LC-RGG1, LC, and RGG1. Standard error of the mean is computed over 3 independent trajectories. The dashed lines are the fitting curves using the following function: R_ij_=b|i-j|. **d** Intramolecular interaction profiles calculated from atomistic simulations of FUS wild type LC-RGG1, LC, and RGG1 single chains, binned by residue position. The left column shows the one-dimensional summation of LC-RGG1 contacts, and the bottom row shows the one-dimensional summation of individual LC and RGG1 contacts. **e** Motions in the condensed phase assessed by ^15^N NMR spin relaxation *R*_1_, *R*_2_, and heteronuclear NOE. Sequence regions of depressed *R*_2_ relaxation are highlighted (grey boxes) and overlayed with possible amyloid forming cores (black boxes). Average relaxation parameters for LC and RGG1 domains in the condensed phase are indicated (grey dashed line) and show that the RGG1 domains show faster reorientational motions than the LC.

To probe the details of interactions of FUS LC-RGG1, we first employed atomistic MD simulations. Previous studies demonstrated a strong correlation between extent of chain collapse of a monomeric protein and the single component phase separation^58,59^. The intrachain distance values show that FUS LC-RGG1 is more collapsed than LC and RGG1 (Fig. 1c), consistent with the enhanced phase separation of LC-RGG1 observed experimentally (Fig 1b).

To identify the regions of the protein chain involved in molecular interactions, we compared the intrachain contacts formed by each independent domain in isolation to the domain in the context of LC-RGG1 (Fig. 1d). We observed that while there are significant heterotypic interdomain-intrachain contacts between LC and RGG1, there are also significant homotypic intradomain-intrachain contacts. Furthermore, the total number and identity of intradomain-intrachain contacts are largely preserved in the LC-RGG1 compared to each isolated domain, although quantitatively reduced (∼15%), as they are exchanged for interdomain contacts. Although tyrosine-tyrosine and tyrosine-arginine show high contact probability, tyrosine and arginine both make contacts with many other residue types including glutamine, serine, and glycine – while other contacts form between pairs excluding tyrosine and arginine, for example glutamine to glutamine (Fig. S1a).

To directly interrogate the interactions that promote LC-RGG1 phase separation, we first experimentally characterized the condensed phase of LC-RGG1. Macroscopic condensed phases for NMR experiments show the same NMR spectral fingerprints as spontaneously formed droplets (Fig. S1b), suggesting these are excellent models for studying phase separated LC-RGG1 consistent with previous work on LC^32^. The condensed phase concentration is approximately 300 mg/ml FUS LC-RGG1 as estimated by NMR intensity referenced to a low concentration reference (see methods). To investigate the molecular motions of LC-RGG1 on the nanosecond timescale, we compared the ^15^N NMR spin relaxation values of LC-RGG1 in the dispersed and condensed phases (Fig. 1e). Overall, the values are consistent with a disordered chain undergoing slower motion in the condensed phase compared to the dispersed phase^60^. Notably, we observed that in the condensed phase, the RGG1 domain exhibits relaxation values consistent with faster reorientational motions compared to those of the LC domain, which is especially clear in the transverse relaxation, *R*_2_, values. This increase in molecular motions may be due to the enrichment of glycine residues in RGG1 which enhance backbone flexibility. We also observed evidence for faster motions in FUS LC between residues 76-96 and 106-121, which have high sequence similarity, are enriched in serine residues, and coincide with segments within the amyloid forming cores of FUS LC^61^.

### Extensive contacts formed involving polar residues in LC-RGG1 condensate

We sought next to probe the domain- and residue-level interactions in the condensed phase of LC-RGG1. To identify how the interactions stabilizing the condensed phase were distributed across the LC-RGG1 sequence, we mixed double labeled (^13^C,^15^N) LC-RGG1 with unlabeled (natural abundance) LC-RGG1 and used filtered/^13^C-edited NOE experiments to characterize the interchain sidechain-sidechain and sidechain-backbone intermolecular interactions (Fig. 2a). NOE intensities were quantified and natural abundance artifacts were removed (see Methods). Consistent with their established role in phase separation, prominent NOEs to tyrosine and arginine positions are observed among the strongest (Fig. 2b, S2a), although complex motions in dynamic phases complicate direct quantification of contact frequency from NOEs^60,62^. Yet, these prominent NOEs are found not only amongst themselves (Tyr-Tyr and Tyr-Arg only, as envisioned in the current molecular grammar) but with other residues as well, especially involving pairs with glutamine that makes up ∼20% of the total sequence composition (Fig. 1a). Furthermore, arginine-arginine NOEs are seen both in this experiment and in filtered/^15^N-edited NOE experiments, suggesting direct RGG1-RGG1 interactions (Fig. 2b,c, S2a,b) may explain the contribution of RGG1 to enhancing and changing the salt dependence of phase separation (Fig. 1b, Fig S1c). Although the RGG1 domain does not readily phase separate on its own under the conditions tested (Fig. S1d), these data suggest RGG1 forms interactions with both LC and RGG1.

**Figure 2.**
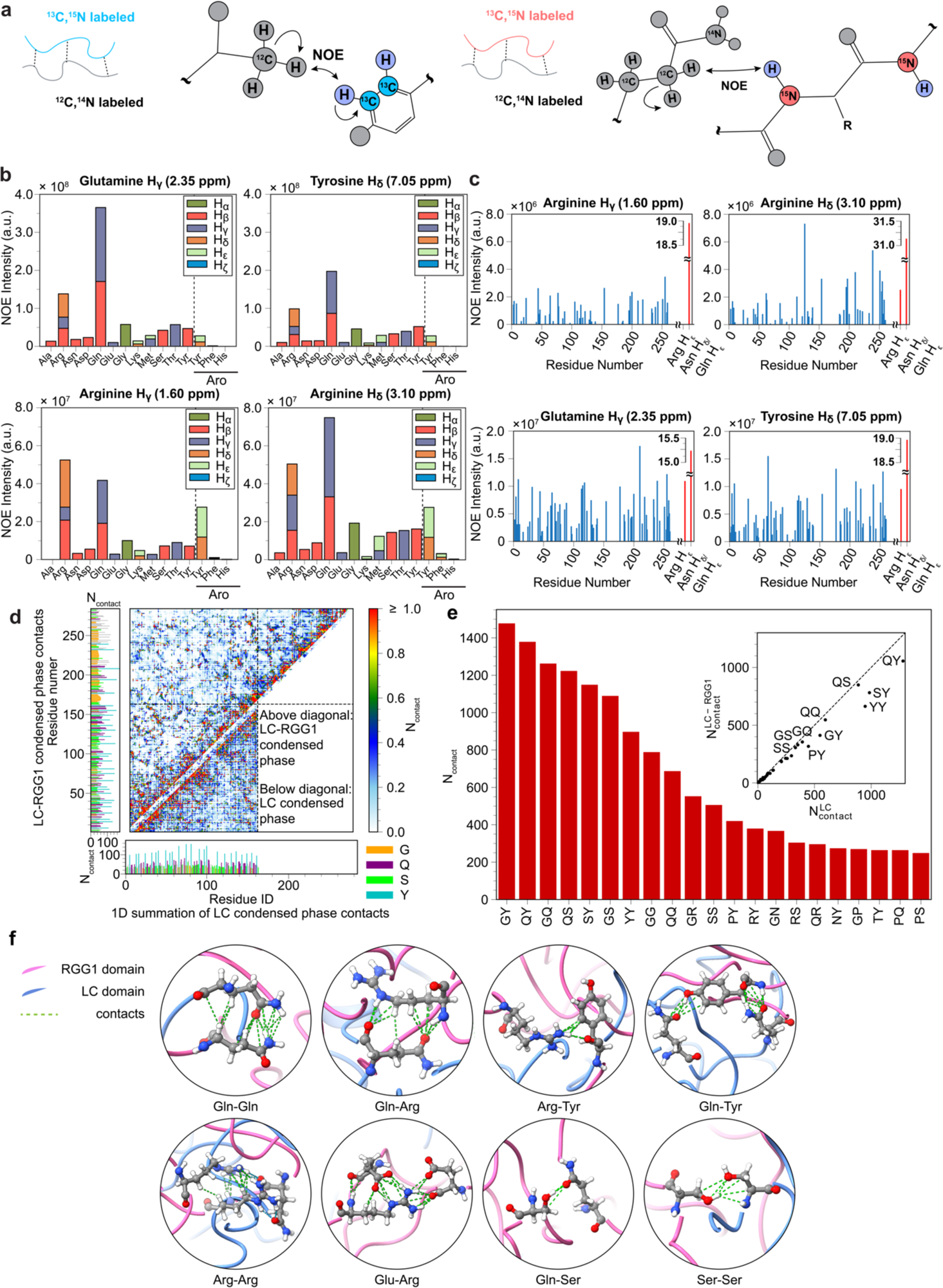
LC-RGG1 makes both homotypic and heterotypic contacts in the LC-RGG1 phase, including aromatic and arginine residues as well as polar residues. **a** Schematic of the ^13^C-(*left*) and ^15^N-(*right*) edited (double ^13^C/^15^N) filtered NOE-based NMR experiments used to measure only intermolecular contacts formed in the LC-RGG1 condensed phase. **b** ^13^C-edited intermolecular NOEs in the LC-RGG1 phase show many residue types have contacts, including LC:LC, LC:RGG1, and RGG1:RGG1 (see Fig. S2a for complete set of observed NOEs). NOE intensities are corrected for (small) intramolecular contributions by subtracting intensities measured in natural abundance control samples (see Methods), and presented as stacked bars for different resolved positions in each residue type. NOEs to aromatic residue positions are measured in a separate experiment and intensities cannot directly be compared to other positions so are presented separated by a dashed line (see Methods). **c** ^15^N-edited intermolecular NOEs show that Gln, Tyr, and Arg intermolecular contacts span across the entire LC-RGG1 sequence, including to side chain Arg and Gln/Asn positions. (see Fig. S2b for complete set of observed NOEs) **d** Interaction profiles calculated from atomistic simulations of FUS wild type LC-RGG1 and LC condensed phase (slab simulations), binned by residue position. The left column shows the one-dimensional summation of LC-RGG1 contacts, and the bottom row shows the one-dimensional summation of LC contacts. **e** The normalized numbers of contacts (by numbers of residues) binned by residue type, calculated from atomistic simulations of FUS wild type LC-RGG1 condensed phase. Inset: Correlation between the pairwise contacts in the LC condensed phased compared to the LC-RGG1 condensed phase. **f** Molecular images of the LC and RGG1 contacts formed in the LC-RGG1 condensed phase simulations, highlighting the different types of residue pair contacts formed in the phase.

To probe the balance of homotypic and heterotypic domain interactions that could be formed, we performed atomistic molecular dynamics simulations of the condensed phase of LC-RGG1 (see Methods for details)^63^, which showed interchain contacts form not only between LC and RGG1 but also between the same domain type (LC and LC, RGG1 and RGG1) (Fig. 2d). The analysis of interchain distances reveals expanded conformations, suggesting a shift from intramolecular to intermolecular interactions. Notably, compared to LC whose conformational properties reflect the expected ideal-chain behavior^64^, the more compact conformations, potentially due to heterotypic interactions between LC and RGG1 (Fig 1c,d). Interactions between LC domains in the LC-RGG1 phase are similar to those formed in the condensed phase of LC alone (Fig. 2d, Fig. 2e inset), as seen for intradomain-intrachain interactions in the dispersed phase (Fig. 1d). Consistent with its dominant role in phase separation of many prion-like proteins including FUS^65,66^, tyrosine makes many contacts (Fig. 2e, S2c). However, these are not uniquely tyrosine-tyrosine and tyrosine-arginine contacts but also with many of the residue types enriched in LC and RGG1. Similarly, arginine also makes prominent contacts with tyrosine but also glycine, glutamine, and serine, consistent with the NMR data. These contacts are stabilized by many interaction modes including hydrogen bonding and hydrophobic interactions (Fig. 2f), in addition to π-π and cation-π interactions^33^. Residues such as Tyr and Arg can interact via many modes simultaneously^32,33,37,63^, which may contribute to their elevated role in phase separation. Specifically, the arginine-arginine contacts we observed by NOE experiments may be stabilized by arginine guanidino groups stacking^67^, combined with additional charge-neutralization electrostatic interactions mediated by negatively charged groups of nearby residues, such as glutamic acid and aspartic acid sidechains (Fig. S3).

### Polar residues make direct intermolecular contacts in the dispersed phase

The condensed phase is of high protein concentration and hence it is possible that the NOEs observed in the condensed phase report not only on contacts that drive phase separation but also on “incidental” interactions that do not appreciably contribute to phase separation^24,68^. Conversely, observation of intramolecular NOEs between non-adjacent tyrosine and phenylalanine positions in a disordered prion-like domain has been suggested to be more stringently controlled evidence for their participation in the driving forces for phase separation as interaction in the dispersed phase would not be incidental^24^. However, in that study only contacts between phenylalanine and tyrosine were probed because essentially all other residue pairs always occur at adjacent positions in these degenerate prion-like domains, preventing analysis of other intramolecular pair contacts. Therefore, we instead used <u>inter</u>molecular NMR experiments to test if the contacts with non-tyrosine polar residues in the FUS LC observed in both NMR experiments and MD simulations are actually observed before phase separation. We chose filtered/edited NOE NMR experiments to rule out any contribution from local sequence (i to i+1, i to i+2, i to i+3, etc.) contacts. Considering the relatively low solubility of FUS LC-RGG1, we chose the LC as a model as it has a higher *C*_sat_ and we further raised *C*_sat_ to create samples at 2.5 mM (10x less concentrated than the condensed phase^32^) by replacing eight tyrosine residues (1/3 of the total) with serine (Fig. 3a). As in condensed phase experiments^32^, we observed NOEs arising from contacts between tyrosine and glutamine as well as glycine and/or serine (Fig. 3b), above any background that may arise from isotope incorporation/abundance or filtering artifacts (we performed the same experiments on both fully labeled and natural abundance control samples, Fig. 3b). To further confirm tyrosine to glutamine contacts in the dispersed phase, we performed intramolecular NOE experiments on a version of FUS where the few YQ or QY adjacent residues are removed (analogous to what was done for F-Y pairs^24^). In these experiments we see tyrosine-glutamine contacts that cannot arise from adjacent positions (Fig. 3b). Therefore, these results of NOEs in the dispersed phase between tyrosine and many other residue types in FUS LC show that the large array of residue-pair NOEs observed in the condensed phase is not an artifact of the high-density and underscores the possibility that intermolecular interactions in the condensed phase are stabilized by contacts with non-tyrosine polar residues. This view is strongly supported by the simulations showing that contacts in the dispersed and condensed phases involving LC to LC, RGG1 to RGG1, and LC to RGG1 domains are highly correlated (Fig. 3c), consistent with recent studies on other proteins^36,37^.

**Figure 3.**
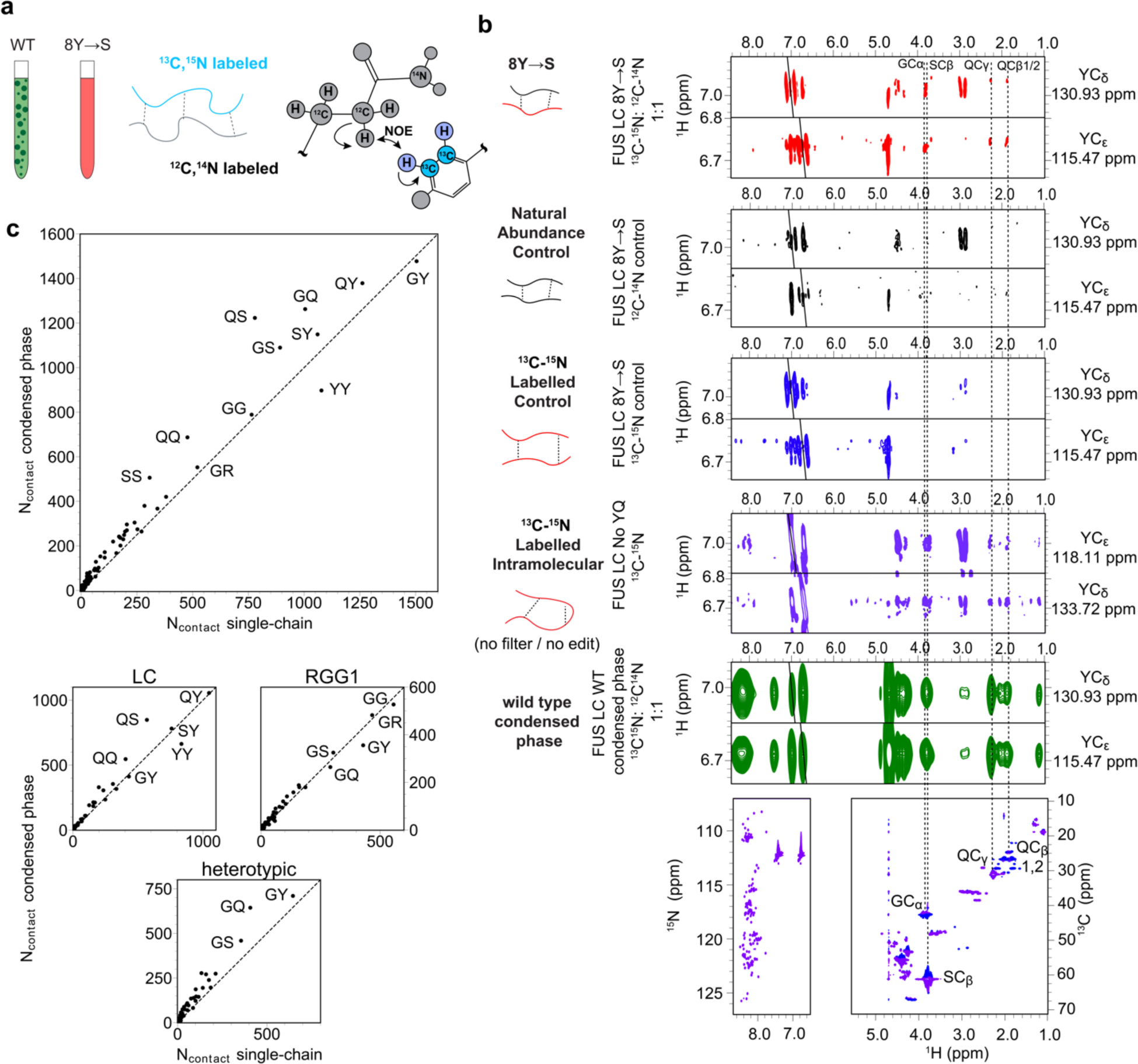
Contacts between tyrosine residues and other polar residues are present in the dispersed phase as well as the condensed phase. **a** Schematic of the samples utilized for intermolecular aromatic ^13^C-edited filtered NOE NMR experiments in the dispersed phase for high concentration FUS LC 8Y→S samples. **b** Using a phase separation deficient FUS LC variant, 8Y→S, intermolecular NOE experiments demonstrate contacts formed between Tyrosine and Glutamine (red, see dashed lines), which are also observed in the condensed phase of FUS LC wild type (green). Using a FUS LC mutant in which i+1/-1 Y/Q sequence pairs are removed, we see that these same interaction are observed in a ^13^C-edited intramolecular NOE experiment for intramolecular contacts. Control samples made with 100% ^13^C-^15^N isotopically labeled (blue) and 100% natural abundance (purple) FUS LC 8Y→S were included as controls to confirm the presence of intermolecular contacts in the mixed sample (red) are bona fide intermolecular NOEs. **c** The correlations between the numbers of contacts formed in the single chain (x-axis) and condensed phase (y-axis) simulations, separated into LC homotypic, RGG1 homotypic, and heterotypic contacts.

### Polar residue identity tunes phase separation

Given the evidence for contacts formed by non-tyrosine polar residues in the condensed phase, we explored the contribution of these residue types to FUS phase separation. We used a mutagenic approach to explore the role of residue types enriched in FUS LC in phase separation. We first substituted all instances of one residue with another (Fig. 4a) and assessed *C*_sat_. To examine the role of glutamine on phase separation, we replaced all glutamine residues in the LC domain (in both FUS LC and LC-RGG1 constructs) with alanine, glycine, asparagine, or serine. Similarly, we changed serine to glycine and threonine to serine, to probe the role of the additional methyl group, and to valine, to probe that of the hydroxyl group. In FUS LC, we were not able to express and purify Q→G or Q→N, though these same variants were tractable in FUS LC-RGG1.

**Figure 4.**
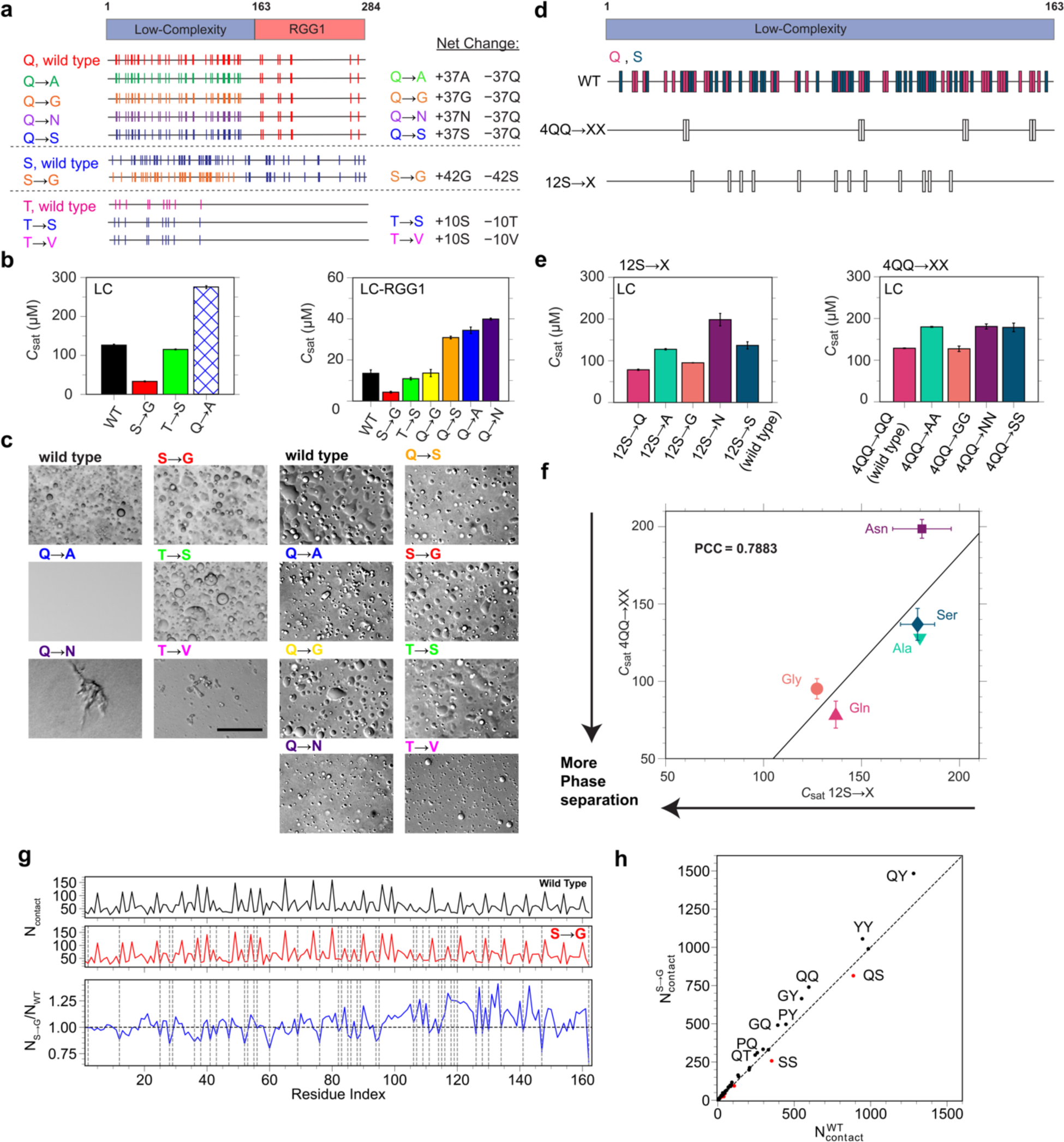
Polar residues alter phase equilibria based on their identity and distribution across the sequence. **a** Schematic of the sequence position of variants used in this study. Complete replacement substitutions were only made in the LC domain of FUS. **b** Measurements of the *C*_sat_ for FUS LC or FUS LC-RGG1 variants in comparison to the wild type. Hatched bar indicates no phase separation. FUS LC experiments were performed at 300 μM and LC-RGG1 at 60 μM in 150 mM NaCl, 20 mM HEPES, pH 7.0 at ambient temperature. **c** DIC micrographs of the substitution variants in the same buffer conditions. **d** Schematic of the FUS LC variants partially modifying polar residues at 4 QQ motifs or 12 S positions. White bars represent which serine or glutamine residues were replaced. **e** Quantification of the effects on the *C*_sat_ of the FUS LC variants vs. the wild type sequence conducted at 300 μM FUS LC in 20 mM HEPES, pH 7.0. **f** Correlation plot between the observed *C*_sat_ for each residue type at 150 mM NaCl condition. **g** 1D profiles of the condensed phase simulation contacts for the wild type and S→G LC mutants, showing higher contacts for residues adjacent to positions where serine to glycine substitutions (dashed lines) were made. **h** Correlation between the number of pairwise contacts formed for the wild type versus the S→G sequence in the simulated condensed phases, showing higher contacts for many pairs including QY and YY in the S→G and fewer contacts for the positions that were converted from serine to glycine (red dots).

We found changes in the phase separation broadly consistent with the prevalence of contacts formed by these residues in stabilizing the condensed phase. First, we see several fold changes in the *C*_sat_ for these substitutions, which do not affect the tyrosine or arginine positions (Fig. 4b, S4a). Second, for the variants that could be performed both in FUS LC as well as FUS LC-RGG1, we see similar effects on the *C*_sat_, suggesting that the presence of RGG1 does not change the contribution of the LC domain to interactions that mediate phase separation. In particular, we see that Q→A, Q→S, and Q→N all similarly decrease phase separation while Q→G is similar to wild-type (Fig. 4b, S4a), implying that both glutamine and glycine have a distinct role compared to alanine, serine, and asparagine. Consistent with these observations and as observed previously for hnRNPA1 LC and A-IDP, S→G enhances phase separation^25,29^. Interestingly, T→S slightly enhances phase separation, suggesting that the methyl group of threonine does not substantially contribute to phase separation. Third, some polar residue substitutions result in aggregation; Q→N in the FUS LC resulted in irregularly shaped aggregates while the same substitution in the LC-RGG1 context remained liquid-like (Fig. 4c). For T→V, the LC variant aggregated immediately while the LC-RGG1 sequence was initially liquid-like but changed to irregular aggregates over the course of approximately 5 minutes (Fig. S4b). Fourth, some substitutions appeared to change the shape of the phase diagram as a function of salt (Fig S4a). In both the LC and LC-RGG1, Q→A is deficient in phase separation except at very high salt concentrations (1M NaCl, Fig. S4c) where the *C*_sat_ for liquid droplets approaches that of the wild-type. These findings as a function of salt suggest the contribution of hydrophobic interactions to phase separation is enhanced at high salt with the more hydrophobic alanine-containing sequence^32,40,69^. Together, these results suggest that non-tyrosine polar residue identity plays an important role in FUS phase separation and aggregation.

Next, to test further how polar residue substitutions alter phase separation without the impact of aggregation and directly compare the different residue types in the same sequence context, we generated sequences where residues were changed at two defined position subsets in FUS LC (Fig. 4d). Expanding on our previous designs^32^, we picked one position subset with four native QQ motifs (4QQ→XX) and the other with 12 native serines (12S→X) and generated constructs that changed these to every other residue type in the following list: glutamine, glycine, asparagine, serine, and alanine. These residues are all found in FUS LC but to differing extents and all are of different sizes and chemical features, allowing comparison of hydrophobic contributions from acyl chain regions, side chain hydrogen bonding, or solvation volume changes on phase separation. First, we find that the polar residue substitutions have overall similar effects in both contexts. Glutamine and glycine substitutions show liquid-like droplet formation and approximately the same *C*_sat_ in the 12S→X and 4QQ→XX contexts (Fig. 4e, S4d,e), like what we observed for replacing all the glutamine with glycine (Q→G) for FUS LC-RGG1. Serine and alanine showed similar reduction of phase separation in both sequence contexts. Interestingly, asparagine showed further reduction of *C*_sat_ compared to serine and alanine in the 12S→X context but similar phase separation in the 4QQ→XX context. This context dependence is clearly seen in the deviation from the correlation plot for these sets of sequences for asparagine (Fig. 4f), suggesting that the contribution of polar residues in phase separation is context/sequence dependent like seen for tyrosine and arginine previously^25,31,36^. Hence, polar residues tune the quantitative details of phase separation, with glutamine and glycine favoring phase separation more than alanine, serine, and asparagine.

### Mechanistic dissection of the role of glycine

Precisely why glycine favors phase separation more than alanine, serine, and asparagine requires more in-depth investigation. Previous interpretations suggested that solvation volume changes, where the residues with no enthalpically favorable interactions to phase separation take up less space when changed to glycine, explain glycine favoring phase separation^25^. However, as we showed above, even removing large portions of side chains with alanine substitutions does not enhance phase separation. Therefore, the effect is likely to be linked to something particular about glycine, perhaps its backbone flexibility, as the distinguishing feature compared to other residues. To gain insight into the role of glycine, we performed atomistic MD simulations of the FUS LC wild type and S→G variant. Compared to wild type simulations, the number of contacts formed by residues adjacent to the mutated position increased in the S→G variant condensed phase simulation (Fig. 4g), which is consistent with the experimentally observed decrease in *C*_sat_ for the S→G variant. Correlation plots comparing the two variants show that QY and YY contacts are increased (along with other residue pair types) while, due to the lack of sidechains in glycine, contacts decreased between the mutated positions (S in the unmutated, G in the mutated - red dots) with other residues (Fig. 4h, S4f). These results suggest that glycine substitution plays an indirect role in phase separation by providing flexibility and allowing for the formation of contacts between other residues that stabilize phase separation, as opposed to enhanced direct glycine interactions (upon serine-to-glycine mutations) with other residues^70,71^

### Liquid-to-solid transition is dependent on non-tyrosine polar residue type

The molecular grammar for the conversion of liquid-like forms to aggregated solids has been summarized in simple rules such as “glycine maintains liquidity, whereas glutamine and serine promote hardening”^6^. To further investigate how the identity of polar residues affects LST, we tested the ability of FUS variants to aggregate as a function of time. We used both quiescent conditions and shaking that generates shear and disruption at phase interfaces that can nucleate aggregation^72,73^, monitoring the onset and morphology of the aggregation over a 24 hour time course using DIC microscopy (Fig. 5a, left). To quantify the extent of amyloid-like fibril formation, we also measured ThT fluorescence over the same time duration (Fig. 5a, right), here with agitation in a plate reader. At quiescent conditions, the FUS LC wild-type, S→G, and T→S all formed round droplets and did not show increases in ThT fluorescence (Fig. 5b,c, S5), consistent with no aggregation. Interestingly, in the LC-RGG1 experiments, the WT forms a few irregular aggregates by the end of the 24-hour period and shows weak ThT enhancement after 4 hours, while neither the S→G nor T→S shows ThT enhancement nor formation of irregular structures by microscopy (Fig. 5d,e, S6). Hence, threonines, whose branched structure at the C_β_ position biases protein chains to form β-sheet structures^74,75^, and serines in the LC domain contribute to FUS LC-RGG1 conversion from liquid-like droplets to solid-like, ThT-positive aggregates. By contrast, LC Q→A does not form droplets or apparent aggregates on the micron-scale but does show weak ThT-positive aggregation after >15 hours of incubation. In LC-RGG1, Q→A and Q→S show similar ThT enhancements that are higher than wild-type and form irregular aggregates by microscopy after several hours, whereas Q→N forms aggregates faster than wild-type and Q→G shows lower ThT enhancement. Hence, although glutamine has been associated with solidification^6^, it is not necessary for prion-like domain aggregation.

**Figure 5.**
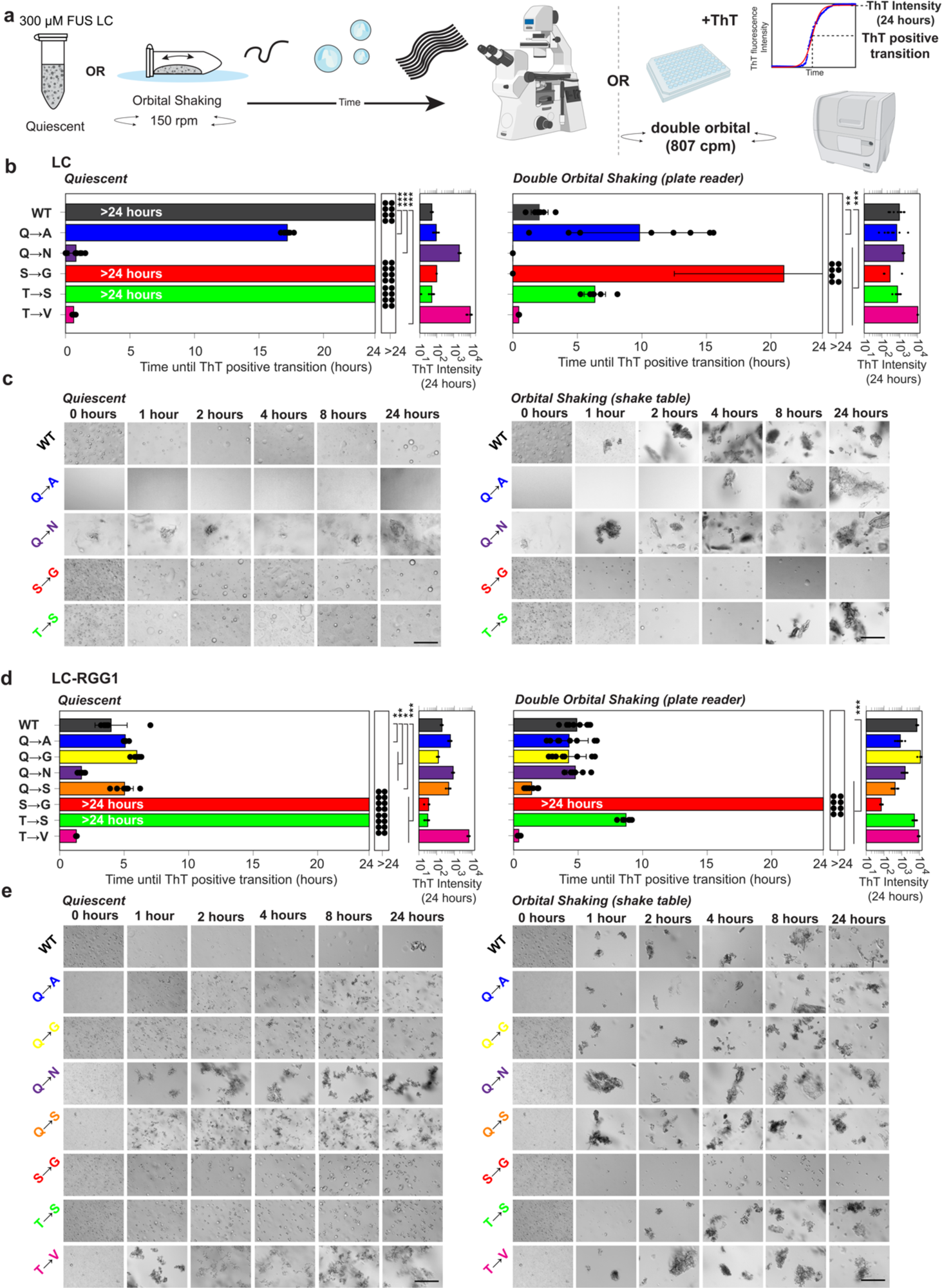
Polar residue identity contributes to the liquid-to-solid transition. **a** Schematics depicting the method of inducing aggregation for the microscopy-based experiment and the ThT based assay. ThT enhancement curves (blue) are fit with a sigmoid (red) and both the ThT enhancement transition and maximum ThT intensity are determined from the fit. **b** Time until ThT positive transition (longer bars indicate ThT-detected aggregation) and ThT intensity at 24 hours (larger bars indicate more ThT enhancement) of the variants in the LC domain when exposed to either quiescent and double orbital shaking conditions. **c** DIC micrographs of the variants in the LC and LC-RGG1 when subjected to 24 hours of quiescent and orbital shaking conditions (150 rpm, up to 24 hours) **d** Same as in **b** but for the variants in the LC-RGG1 sequence (24 hour duration, double orbital shaking). **e** Same as in **c** but for the variants in the LC-RGG1 sequence under quiescent or 150 rpm shaking conditions and up to 24 hours.

We also performed assays with rapid shaking to compare the variants further. Here we see similar results, with the main difference being that aggregation proceeds faster in most cases with shaking. With shaking, some sequence-specific differences are also evident. T→S, which was not ThT positive and did not form aggregates at any timepoint at quiescent conditions, does form aggregates detected by microscopy and ThT enhancement at approximately 8 hours. Hence, the native threonine positions are not required for aggregation but do speed aggregation. Even at these harsh shaking conditions, S→G remains liquid-like and does not show ThT positivity. Given that Q→G forms aggregates but S→G does not, these data suggest that serines, but not glutamines, are required for FUS LC-RGG1 aggregation.

### Sequence-specific impact of polar residue identity on aggregation points to specific amyloid core formation

Based on these observations, we used our two distinct subset variants (12S→X and 4QQ→XX) to test the sequence-specific impact of polar residue identity on aggregation – specifically, what role serine has at these positions. We subjected these LC sequences to the same aggregation assays as above. As expected, based on replacement of all glutamines with other polar residues (Fig. 6a,b, S7), substitutions of only some glutamines (4QQ→XX) only slightly changed the behavior compared to the wild type for alanine, serine, and glycine – the droplets remained spherical at long times and no irregular shaped aggregation was observed at quiescent conditions. Surprisingly, serine substitution (4QQ→SS) showed slower conversion to irregular morphologies by microscopy compared to the wild-type, but this difference was not clear from the ThT assays, suggesting that not all prion-like domain aggregates can be equally detected by amyloid-binding dyes. Similarly, asparagine (4QQ→NN) showed irregular shaped droplets after one hour even at quiescent conditions, showing that glutamine to asparagine substitutions are more prone to aggregation – but these aggregates are not ThT positive (Fig. S8). Strikingly, the other series of subset changes altering native serine positions (12S→X) showed significant effects on aggregation under shaking conditions. We found that the wildtype with native serine was most aggregation-prone while substitutions of serine with alanine, glycine, asparagine or glutamine slowed aggregation based on both morphology and ThT fluorescence.

**Figure 6.**
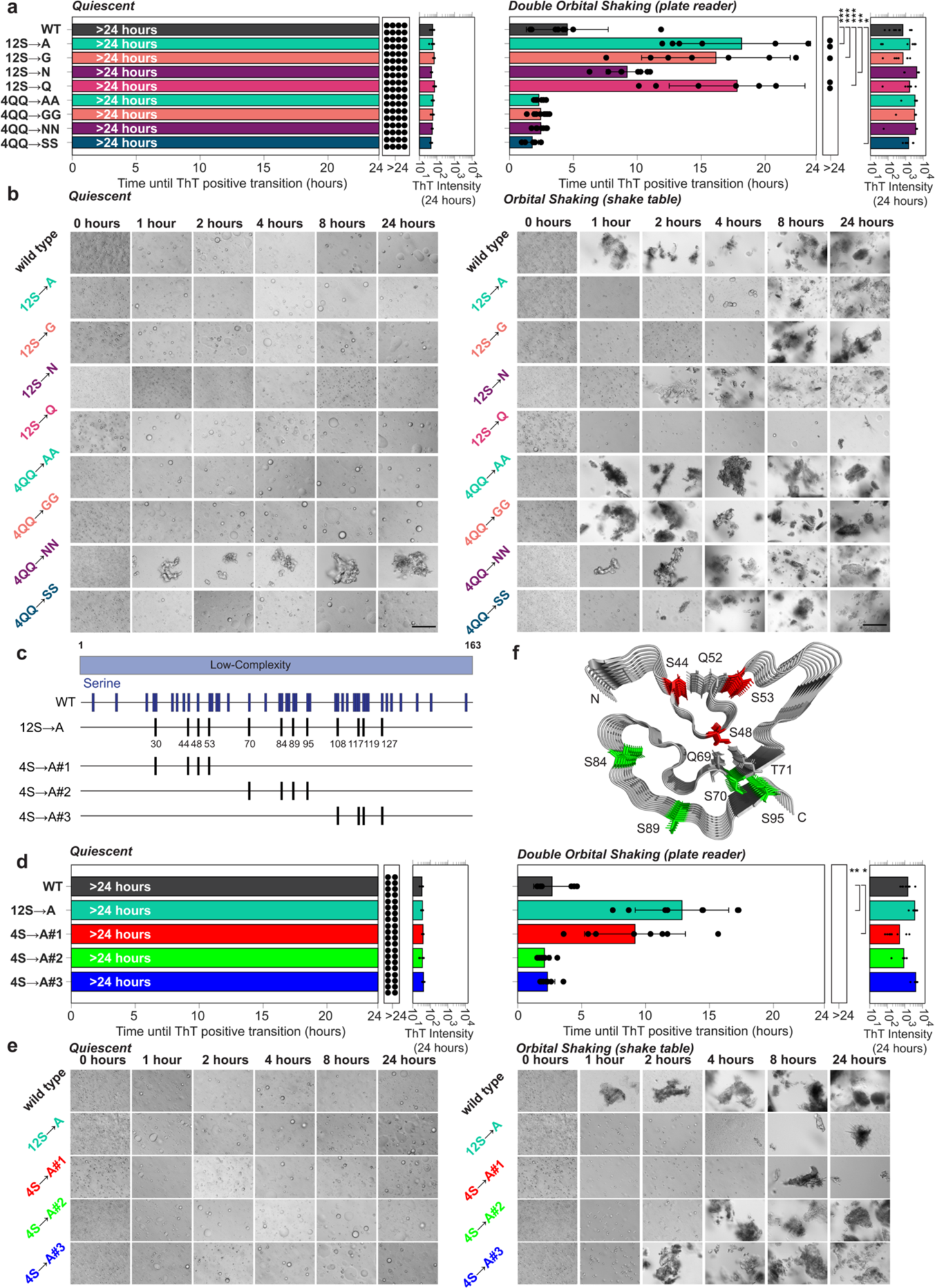
Serine side chain specifically contributes to formation of aggregates from FUS LC droplets. **a** Time until ThT positive transition and ThT fluorescence at 24 hours of FUS LC variants with partial polar residue substitution when subjected to either quiescent or aggregation-inducing conditions (double orbital shaking). **b** DIC micrographs of FUS LC variants with partial polar residue substitution when subjected to quiescent or aggregation-inducing mutations (150 rpm shaking). **c** Schematic of the serine to alanine sequences generated to determine which serines contribute most significantly to aggregation in FUS LC. **d** same as **a** but for the 4S→A variants under quiescent or aggregating conditions (double orbital shaking) to determine which subset of the 12 S→A serines are most important for aggregation **e** same as **b** but for the 4S→A variants under quiescent or aggregating conditions (150 rpm shaking) **f** Serines modified in this study mapped to their location in a proposed amyloid core (PDB:5W3N) for serines in 4S→A#1 (red) and 4S→A#2 (green). Glutamine residues identified as forming hydrogen bonds in the structure (gray sticks) are found in 4S→A#1.

Given that changing only 12 serines could prevent FUS LC aggregation, we then aimed to further narrow the specific sequence regions of FUS LC that contributed the most to aggregation (Fig 6c). Recently, several distinct amyloid-forming cores for FUS LC have been identified by solid-state NMR^76,77^, however, it is unclear which if any of them account for FUS LC aggregation from liquid-like condensed phases. To test which sequence regions were responsible for initiating β-sheet driven aggregation, we created three sequences that replaced only four of the twelve serines with alanine and subjected these peptides to our aggregation assays (Fig. 6d,e, S9). We found that substitutions of the first four positions (S30A/S44A/S48A/S53A) delayed aggregation nearly identically to 12S→A, while the two other variants with four serine to alanine substitutions were statistically indistinguishable from the wild-type. This finding shows that these particular serine residues have critical involvement in aggregation and suggests that the amyloid core reported by Murray *et al.*, which is stabilized by hydrogen bonds between S48 and Q69 and T71 as well as S44 with Q52^77^ may be the structure formed here (Fig. 6f). At the same time, the finding that aggregation is markedly decreased by 12S→A while phase separation is not qualitatively affected suggests again that formation of these β-sheet cores is not required for FUS LC phase separation^32^.

## Discussion

Specifying functional phase separation and avoiding deleterious aggregation are important features for disordered domains. Hence it is important to understand the molecular grammar underlying disordered domain self-assembly and aggregation. Aromatic residues (tyrosine, tryptophan, and phenylalanine depending on the context) have been seen as the ones that form the only dominant contacts with each other (via π-π interactions), while arginine-aromatic contacts (via cation-π interactions) play an auxiliary role – the other residues (Q/S/G/T/N/A) that are enriched in these domains are not thought to form stabilizing contacts or contribute to the molecular driving forces. We set out to test what if anything the polar and small residues contribute to phase separation and what the rules are for their contribution to aggregation.

First, we found a surprisingly richer view for the role of arginine in phase separation than previously proposed as primarily interacting with aromatic residues or oppositely charged residues. By probing the phase separation of FUS LC-RGG1, which combines a tyrosine rich low complexity domain and an RGG domain enriched in charged resides, we find that RGG1-RGG1 and LC-RGG1 contacts contribute to phase separation. In the condensed phase, arginine forms contacts with many residue types including glutamine and other arginine residues.

Second, we found that residue types in FUS other than tyrosine and arginine contribute favorably to phase separation. Here we show that glutamine is an important contributor to phase separation, forming contacts in both the dispersed and condensed phases that contribute to the thermodynamics of phase separation. This holds in several sequence contexts including replacing it in the entire FUS LC and also at a subset of native serine and native glutamine positions. Interestingly, despite the similarity in the functional end group, asparagine substitution for glutamine decreases phase separation, suggesting that glutamine’s aliphatic region makes contributions to phase separation, consistent with hydrophobic contacts at these sites^25,32^ or enhanced length/flexibility near the hydrophilic end group. Furthermore, because asparagine, serine, and alanine all reduce FUS phase separation (as compared to glutamine) but are all smaller than glutamine, the effect of glutamine on phase separation cannot be explained by solvation volume. Importantly, we saw this effect regardless of whether the native position was glutamine or not (i.e. both in 12S→X and 4QQ→XX comparisons). This is in direct contrast to what was seen previously for hnRNPA1, where glutamine substitutions at 14 asparagine positions decreased phase separation^25^. With only a few native asparagine residues, it is difficult to test the role of native asparagine positions in FUS LC. In summary, though making a smaller contribution on a per-residue basis than tyrosine, glutamine in FUS appears to be a driver of phase separation, showing the importance of evaluating the contribution of residue types in different contexts in order to reveal the complete molecular grammar. Indeed, glutamine is prominently featured in prion-like domains and repeat expansions associated with phase separation^78,79^. Hence, the established molecular grammar for phase separation of prion-like domains could be expanded to include glutamine at the very least. Indeed, we find that glutamine enhances phase separation much more than alanine, consistent with our previous results showing that methionine also contributes more than alanine to TDP-43 phase separation^30^. Alternatively, given the role of many different residue pair contacts in phase separation, one could adopt a non-binary inclusive classification where most (if not all) residues can contribute toward molecular driving forces even if primarily via forming stabilizing contacts with other residue types, depending on the context and other environmental factors such as temperature, salt, and pH^80–82^.

Third, glycine plays a special role in phase separation that cannot be ascribed to solvation volume. Here, we show that glycine to glutamine substitutions show little change in phase separation, unlike glutamine substitutions with any other polar/small residue. However, glycine enhances phase separation compared to serine^25^, alanine, and asparagine in all FUS sequence contexts. Molecular simulations suggest that glycine may enhance the contacts formed by other adjacent residues, including tyrosine, by adding flexibility to the backbone. Therefore, the role of glycine in phase separation may be due to its ability to adopt conformations unfavorable for other amino acids.

Fourth, we provide an update to the molecular grammar of phase separation of prion-like domains. The primacy of tyrosine and arginine residues in impacting phase separation in residue substitution experiments has contributed to the view that these residue types function as *sole* drivers of phase separation wherein they have favorable interactions only with themselves. However, in our view, mutagenesis experiments cannot establish pairwise contacts and modes that drive phase separation. NMR experiments and all-atom simulations of condensed phase do not support a view that favorable contacts are only formed between tyrosine and arginine – indeed tyrosine and arginine make up only 15% of the sequence and so ∼1% of the possible pair interactions. Based on our findings here that tyrosine and arginine can form contacts with glutamine and other polar residues in the dispersed phase as well as the condensed phase, we suggest that tyrosine and arginine play a dominant role in phase separation because they form favorable interactions important for stabilizing phase separation with themselves as well as with many residue types. Therefore, we propose viewing disordered domain phase separation through the lens where all residues types may form contacts with a continuum of favorable contributions to phase separation. Importantly, this model is consistent with pioneering work that established the hierarchy of residue-type contribution to phase separation, however here we show that residues like tyrosine and arginine form many stable contacts with many residue types. Support for this view also comes from multiple highly successful coarse-grained (CG) models of polypeptide phase separation^83^, which show a continuum of pair-wise interaction strength, as would be anticipated from the multiple chemical modes of interaction between residues^32^ as well as the high local mobility of residues in the condensed phase^35,84,85^ that suggest contacts are distributed, not localized, and dynamic, not long lived, unlike what would be expected in a model with few localized strong interactions^86^. These distinctions in models matter because the way experiments are designed, and data interpreted needs to be through a physically accurate lens.

Finally, we find that the molecular grammar of polar residue contribution to the solid-to-liquid transition is context and position dependent. Substitutions S→G and T→S that decrease the β-sheet torsion angle propensity of the backbone can decrease aggregation. However, substitutions which only change sidechain details follow much less clear rules. We find that although glutamine was suggested to be important for FUS solid-to-liquid transition, substitutions at all native glutamine positions to glycine, serine, alanine, and asparagine also form ThT-positive assemblies consistent with amyloid-like aggregates. However, asparagine does stabilize aggregation more than glutamine in many sequence contexts^87,88^. For native serine positions, we find that substitutions to glutamine or any other polar/small residue type delayed aggregation, which we found was associated with a subset of positions between residues 30 and 53 in FUS LC, mapping to the beginning of the FUS LC fibril core found by McKnight and coworkers^77^. Hence, aggregation is disrupted by backbone flexibility but is formed via specific side-chain details that defy simple rules, thereby highlighting the need for additional studies to probe aggregation behavior in a variety of sequence contexts, as suggested by the number of disease-associated mutations leading to aggregation of prion-like domains.

Together, these data provide rich insight into the residue-dependence of phase separation and liquid-to-solid transition for FUS, expanding the molecular grammar needed to understand the roles of this family of proteins in physiology and their dysfunction in disease.

## Methods

### Protein Expression and Purification

Recombinant expression and insoluble purification of hexa-Histidine tagged FUS LC and LC-RGG1 wild type (WT) and mutant FUS LC and LC-RGG1 was performed in *E. coli* using previously described methods^32^. FUS LC and LC-RGG1 variants were purified using the same methods as the wild type. To achieve desired isotopic labeling schemes, *E. coli* were grown in M9 media (27 mM NaCl, 22 mM KH_2_PO_4_, 51 mM Na_2_HPO_4_·7H_2_O, 1 mM MgSO_4_, 1%v/v MEM Vitamin Solution (100x), 0.2%v/v solution Q)) using ^13^C-glucose and/or ^15^N-ammonium chloride as the sole carbon or nitrogen source, respectively.

### NMR Sample Preparation and NMR Spectroscopy

NMR experiments were recorded on either a Bruker Avance III HD 850 MHz spectrometer equipped with a HCN TCI cryoprobe with z-gradients, where indicated. All experiments were processed using NMRPipe and CCPNMR 2.5.2.

Condense phase samples of FUS LC-RGG1 were generated by ten fold dilution of concentrated FUS LC-RGG1 wild type (3 mM or greater) from 20 mM CAPS, pH 11.0 into 20 mM MES pH 5.5, 150 mM NaCl, 10% D_2_O and then centrifuged at 4000 rpm at 22 °C for 10 min. The condensed phase was then incrementally transferred to a 3 mm NMR tube which was centrifuged at 1000 rpm for 10 min at a time at 22 °C.

Condensed phase protein concentration was estimated using a one dimensional ^1^H NMR spectrum (zgpr) with an ultra-weak presaturation pulse as previously reported^32^. Data was collected with a center frequency of 4.7 ppm and 32768 time domain points and a spectral width of 15 ppm. The area of the amide backbone envelope between 6.5 – 9.0 ppm was integrated and compared with a low concentration FUS LC-RGG1 dispersed phase standard.

High concentration dilute phase samples of FUS LC 8Y→S were generated by spin concentration in 20 mM CAPS pH 11.0 using a 3 kDa MWCO centrifugal spin filter unit with 0.5mL capacity (Amicon). The sample was then diluted into 50 mM MES pH 5.5, 150 mM NaCl, and 10% D_2_O and subsequently transferred to a 3 mm NMR tube for analysis.

^1^H ^15^N HSQCs for the dilute, biphasic, and condensed phases were acquired with 3072 direct time domain points and 400 indirect points with spectral widths spanning 10.5 and 20 ppm in the direct and indirect dimensions, respectively. The ^1^H and ^15^N carrier frequencies were set to 4.7 and 117 ppm, respectively, and direct and indirect signals were collected for 172 and 116 ms each.

Motions of the FUS LC-RGG1 backbone in the condensed and dilute phase were measured using standard pulse sequences (hsqct1etf3gpsitc3d, hsqct2etf3gpsitc3d, hsqcnoef3gpsi). ^15^N longitudinal relaxation (*R*_1_) was measured using a 7-point interleaved relaxation delay consisting of (100, 1000, 200, 800, 300, 600, 400 ms). Spectra at each delay value were acquired with 4096 direct time domain points and 256 indirect points centered at 4.7 and 117 ppm and with spectral widths of 10.5 and 20 ppm for ^1^H and ^15^N, respectively. ^15^N transverse relaxation (*R*_2_) was measured using a 7-point variable relaxation delay (16.9, 270.4, 185.9, 33.8, 118.3, 84.5, 169 ms) using a 556 Hz CPMG field. Spectra at each point were acquired with 4096 direct time domain points and 256 indirect points, carrier frequencies set at 4.7 and 117 ppm, and spectral widths of 10.5 and 20 ppm, respectively. Time domain data was acquired for 229 ms in the direct dimension and 74 ms in the indirect dimension. ^15^N heteronuclear NOEs were collected using interleaved steady-state NOE and no-NOE control experiments, each with a 5 s recycle delay. Spectra were collected with a center frequency of 4.7 and 117 ppm, spectral widths of 10.5 and 20 ppm, and acquisition times of 172 and 149 ms for ^1^H and ^15^N, respectively.

Intermolecular NOE experiments (noesyhsqcgpwgx13d) exploring sidechain-sidechain interactions were recorded on the condensed phase containing 1:1 mixture of ^13^C/^15^N:^12^/^14^N FUS LC-RGG1 and 1:1 mixture of ^13^C/^15^N:^12^C/^14^N FUS LC 8Y→S. For FUS LC-RGG1 samples, aliphatic and aromatic regions of the spectrum were collected separately. For the aliphatic regions, a ^13^C carrier frequency of 43 ppm was used while the homonuclear ^1^H carrier frequency was set to 4.7 ppm. Spectra were obtained with 4096, 64, and 128 total points spectral widths of 12, 80, and 12 ppm in F3, F2, and F1, respectively. A mixing time of 100 ms and 1 s recycle delay was used. For the aromatic regions, a ^13^C carrier frequency of 80 ppm was used with homonuclear ^1^H carrier frequencies set to 4.7 ppm. The aromatic centered NOE experiments were collected with the same spectral parameters as described above. Control samples with 100% natural abundance FUS LC-RGG1 were created, intramolecular NOE’s arising from natural abundance ^13^C and ^15^N were measured, and NOE intensity values were multiplied by 0.5 and subtracted from the intermolecular NOE data. For FUS LC 8Y→S samples, the ^13^C carrier frequency was set to 122.5 while ^1^H carrier frequency was set to 4.7. These data were collected with 4096, 16, 512 time domain points and spectral widths of 12, 45, and 12 ppm for F3, F2, and F1, respectively. These experiments were collected with a 150 ms mixing time and a 1 s recycle delay.

Intramolecular NOE experiments (noesyhsqcetgp3d) examining self-interaction within the dilute phase were recorded on 150 μM ^13^C/^15^N FUS LC samples in 50 mM MES, pH 5.5, 150 mM sodium chloride, 5% D_2_O. Spectra corresponding to the aliphatic and aromatic regions were collected separately. For the aliphatic region, the ^13^C carrier frequency was set to 40 ppm, while the ^1^H channels were centered at 4.7 ppm. The spectra were acquired with 3072, 64, and 128 total points and spectral widths of 10, 60, and 8.6 ppm for the F3, F2, and F1 dimensions, respectively. A 250 ms mixing time was utilized with a 1.2 s recycle delay. The aromatic regions of the spectrum was obtained by setting the ^13^C carrier frequency to 112 ppm, while both ^1^H carrier frequencies were set to 4.7 ppm. The spectral widths consisted of 10, 44, and 8.6 ppm with total points of 3072, 32, and 256 in F3, F2, and F1, respectively.

Intermolecular NOE experiments (noesyhsqcf3gpwgx13d) exploring sidechain-backbone interactions recorded on the condensed phase containing 1:1 mixture of ^13^C/^15^N:^12^/^14^N FUS LC-RGG1. These data were collected with a ^15^N carrier frequency of 117 ppm and ^1^H carrier frequencies set to 4.7 ppm. Spectra were acquired with 4096, 128, and 128 total points and spectral widths of 12, 20, and 12 ppm for F3, F2, and F1, respectively. A 100 ms mixing time and 0.8 s recycle delay was used. Control samples with 100% natural abundance FUS LC-RGG1 were created, intramolecular NOE’s arising from natural abundance ^13^C and ^15^N were measured, and NOE intensity values were multiplied by 0.5 and subtracted from the intermolecular NOE data.

### Phase Separation Quantification

The salt dependent phase boundary was quantified using commonly utilized methods^32^. Determination of the FUS LC and LC-RGG1 phase boundaries were conducted using either 300 or 60 μM LC or LC-RGG1, respectively. These experiments were conducted by ten fold dilution of the protein into 20 mM HEPES pH 7.0 and increasing sodium chloride concentrations (0, 75, 150, 300, 600, 1000 mM) from an initial solution of 20 mM CAPS, pH 11.0. After preparation, these samples were centrifuged for 20 minutes at 14,000g at 24 °C. The supernatant was then sampled and its absorbance at 280 nm measured to determine the concentration of the protein remaining in the supernatant.

### Turbidity Measurements

Turbidity measurements of FUS LC, LC-RGG1, RGG1, and mixtures of FUS LC and RGG1 at varying concentrations were conducted in 20 mM HEPES buffer at pH 7.0 in the presence of 150 mM sodium chloride and at 25°C. The absorbance at 600 nm was obtained using a Cytation 5 plate reader. Individual samples were prepared and 90 μL of each were transferred to a 96-well plate (Costar) and sealed with clear optical adhesive.

### Microscopy

Evaluation of droplet morphology for each construct and the microscopy-based aggregation assay was obtained using a Nikon Ti2 DIC microscope with 20x objective and 1.5x digital enhancement for all images. For evaluation of droplet morphology, samples were generated in 20 mM HEPES, pH 7.0, and 150 mM sodium chloride and 20 μL was transferred to a glass cover slip for imaging.

Images obtained to demonstrate that the 4NN sequence was not responsive to ThT under quiescent conditions, brightfield images were obtained using a Cytation 5 plate reader equipped with 10x objective. ThT fluorescence measurements were then obtained on the same sample using the method listed below.

### DIC Microscopy-based Aggregation Assay

Samples containing either 300 or 60 μM protein for FUS LC and LC-RGG1, respectively, were prepared in 20 mM HEPES, pH 7.0 with 150 mM sodium chloride. Independent samples were generated for each time point (0, 1, 2, 4, 8, and 24 hours). A sample volume of 200 μL in 1.5 mL Eppendorf tubes was necessary to prevent the liquid from becoming retained in the tip of the tube. The samples were placed into an Infors HT Multitron Standard incubator capable of 150 rpm orbital shaking and fixed horizontally such that the liquid bead was able to migrate along the length of the sideways Eppendorf tube, without becoming trapped at either end. The temperature was held constant at 25 °C. At the end of each time point, samples were thoroughly mixed and 20 μL of each sample was transferred to a glass cover slip for imaging. Quiescent conditions were achieved by independently preparing the same sample scheme and allowing the samples to sit undisturbed on the benchtop and imaged similarly at the end of each respective time point.

### Thioflavin T Fluorescence Assay

To quantify the aggregation of FUS LC and LC-RGG1 constructs, Thioflavin T (ThT) fluorescence was measured using a Cytation 5 plate reader. Eight samples of either 300 or 60 μM FUS LC or LC-RGG1, respectively, were generated in 20 mM HEPES, pH 7.0, 150 mM sodium chloride and 20 μM ThT. Next, 90 μL of each sample was mixed well and then transferred to a 96-well plate. Double-orbital shaking at 807 cpm at 25 °C or quiescent conditions were utilized. The ThT response curves were then used to best fit a sigmoid form according to the equation, y=d+a/(1+exp(-b*(x-c))), and the inflection point was used to determine the representative aggregation time for each replicate to reach ThT positivity. The maximum ThT fluorescence at 24 hours was obtained from this fit procedure. Fitting parameters were bounded by experimentally relevant values (0<t<24 hours) and background ThT fluorescence was accounted for by measuring ThT fluorescence in the absence of protein. It was necessary to set a threshold requirement that the maximum fluorescence enhancement value must be at least 1.2x above the initial value to prevent low-fidelity fitting of time series where no fluorescence enhancement was observed.

### General Protocol of Atomistic Molecular Dynamics Simulations

Initial equilibration simulations were performed using the classical MD package GROMACS-2022^89^, employing periodic boundary conditions and a 2 fs integration time step. All systems were simulated in explicit solvent, modeled using the AMBER03ws force field^90^ and TIP4P/2005s water model^90^ with improved salt (NaCl) parameters^91^. The initial protein structure was solvated with counter ions added to achieve electroneutrality and a salt concentration of 100 mM. The solvated protein system then underwent energy minimization with the steepest descent algorithm, followed by a 100 ps NVT equilibration. Velocity rescaling algorithm was used for temperature equilibration at 300 K with a coupling constant of 0.1 ps. Subsequently, a 100 ps NPT equilibration was conducted using the Parrinello-Rahman barostat^92^ with a coupling constant of 2 ps for pressure control.

The production runs were performed with MD package Amber22^93^. Post GROMACS equilibration, the package ParmEd^94^ was used to convert GROMACS files to Amber files. Hydrogen mass repartitioning^95^ was applied during conversion to enable a timestep of 4 fs for production runs. Following the conversion, minimization was applied with protein position restraint, followed by a 2 ns NVT equilibration with reduced protein position restraint. With Berendsen barostat^96^, a 500 ps NPT equilibration with further reduced protein position restraint was conducted. After the equilibration steps, the production run simulations were performed in the NVT ensemble. The Langevin dynamics were used to control temperature at 300 K (friction coefficient = 1 ps⁻¹), and the SHAKE algorithm, as implemented in Amber22, was used for constraining hydrogen-containing bonds.

After production runs, the Amber NetCDF trajectory files were converted to Gromacs compressed trajectory files via CPPTRAJ^97^ package for further analysis.

### Initial Structure of All-atom Single Chain Simulations

The initial conformations of single chain simulations are generated from our CG–HPS model^20,21^. Selecting 3 conformations with its radius of gyration (R_g_) was around the mean R_g_ (<R_g_>), < R_g_ > plus 1 standard deviation (std), and < R_g_ > minus 1 std, over 1 μs CG simulation. All-atom structures are reconstructed from these 3 conformations with the package Modeller^98^. The backmapped all-atom structures then proceeded to the equilibration and production simulations, following with the above protocol.

### Initial structure of all-atom slab simulations

Based on the protocol described before^63^, the CG dense phase configuration in a slab geometry was used to reconstruct all-atom slab configuration with Modeller^98^. Any conflicts between side chains of were resolved via a short simulation with the OpenMM-7.6 python package^99^ and Amber99sb-ildn force field^100^ and corresponding implicit solvent model^101^. The relaxed protein system proceeded to equilibration simulations and further production simulations using the protocol described above.

### Initial Structure of FUS LC S→G Variant Condensed Phase Simulation

These simulations started from the equilibrated wild-type protein structure, used “swapaa” command in software Chimera^101^ to replace serine residues to glycine. After removing any atomic conflicts with OpenMM, the resulting structure was subject to equilibration and further production simulation.

### Contact Map Calculation

The package MDAnalysis-2.5.0^102,103^ was used to calculate pairwise residue contacts. Contacts were considered formed if any heavy atom from each of the two residues are within 6 Å distance. Residue pairwise contacts are computed by summation of all the contacts formed by each heavy atom pair between the residues. Except these contact definitions specified here, the analyses are the same with our previous work^63^.

## Acknowledgements

We thank Dr. Mandar Naik for NMR assistance and the Structural Biology Core Facility at Brown University. This work was primarily supported by National Institutes of Health grant R01GM147677. Additionally, the work was partly funded by National Science Foundation (NSF) grant 1845734 (N.L.F.) and Welch Foundation A-2113-20220331 (J.M.). N.W. was supported in part by an NIGMS training grant at Brown University (T32GM139793). T.Z. was supported in part by a Pape Adams Postdoctoral Award from the Carney Institute for Brain Science at Brown University. We gratefully acknowledge the computational resources provided by the Texas A&M High Performance Research Computing (HPRC). This content solely reflects the authors and does not necessarily represent the official views of the funding agencies.

