## Supplemental Figures for "Expanding the molecular grammar of polar residues and arginine in FUS prion-like domain phase separation and aggregation"

### Supplementary Figures

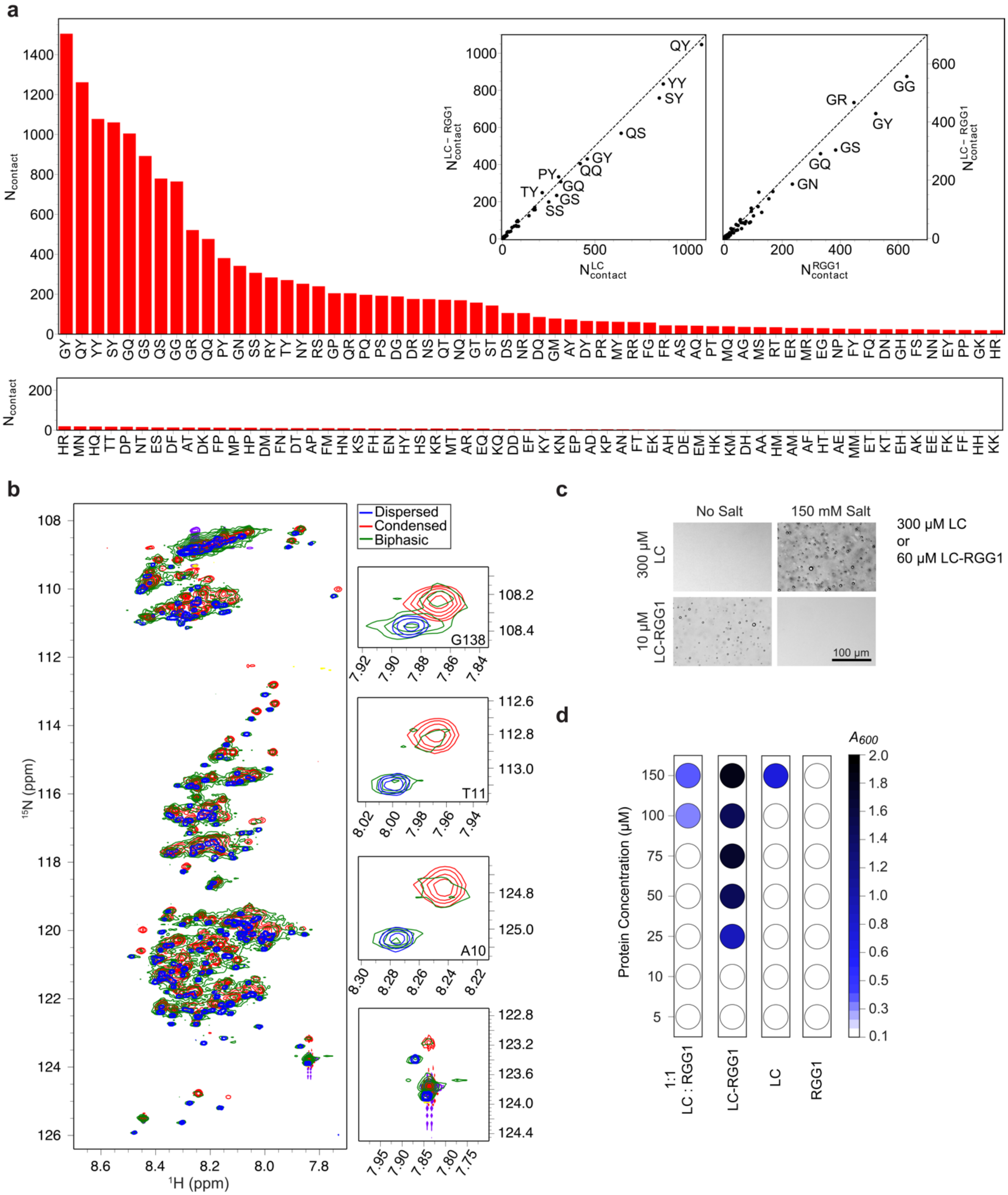


**Figure S1. Phase separation and dilute phase contacts formed by FUS LC-RGG1**

**a** Pairwise contacts formed in the isolated monomer (single chain) simulations of FUS LC-RGG1. Inset: correlation plots of the number of pair contacts formed by the isolated FUS LC (left) or FUS RGG1 (right) domains compared to that domain in FUS LC-RGG1.

**b** $^{1}H$-$^{15}N$ HSQC spectra of FUS LC-RGG1 biphasic sample (green) showing resonances corresponding to both dispersed (blue) and condensed (red) phases at the same time, as represented in the insets.
**c** DIC micrographs of FUS LC (300 μM) or LC-RGG1 (60 μM) in 20 mM HEPES pH 7.0 in the presence and absence of 150 mM sodium chloride, showing the salt dependence of phase separation.
**d** $A_{600}$ turbidity measurements of FUS LC, RGG1, LC-RGG1, and a stoichiometrically equivalent mixtures of LC + RGG1, showing that FUS LC-RGG1 phase separates more avidly than FUS LC, FUS RGG1 or the "in *trans*" mixture of FUS LC and FUS RGG1.

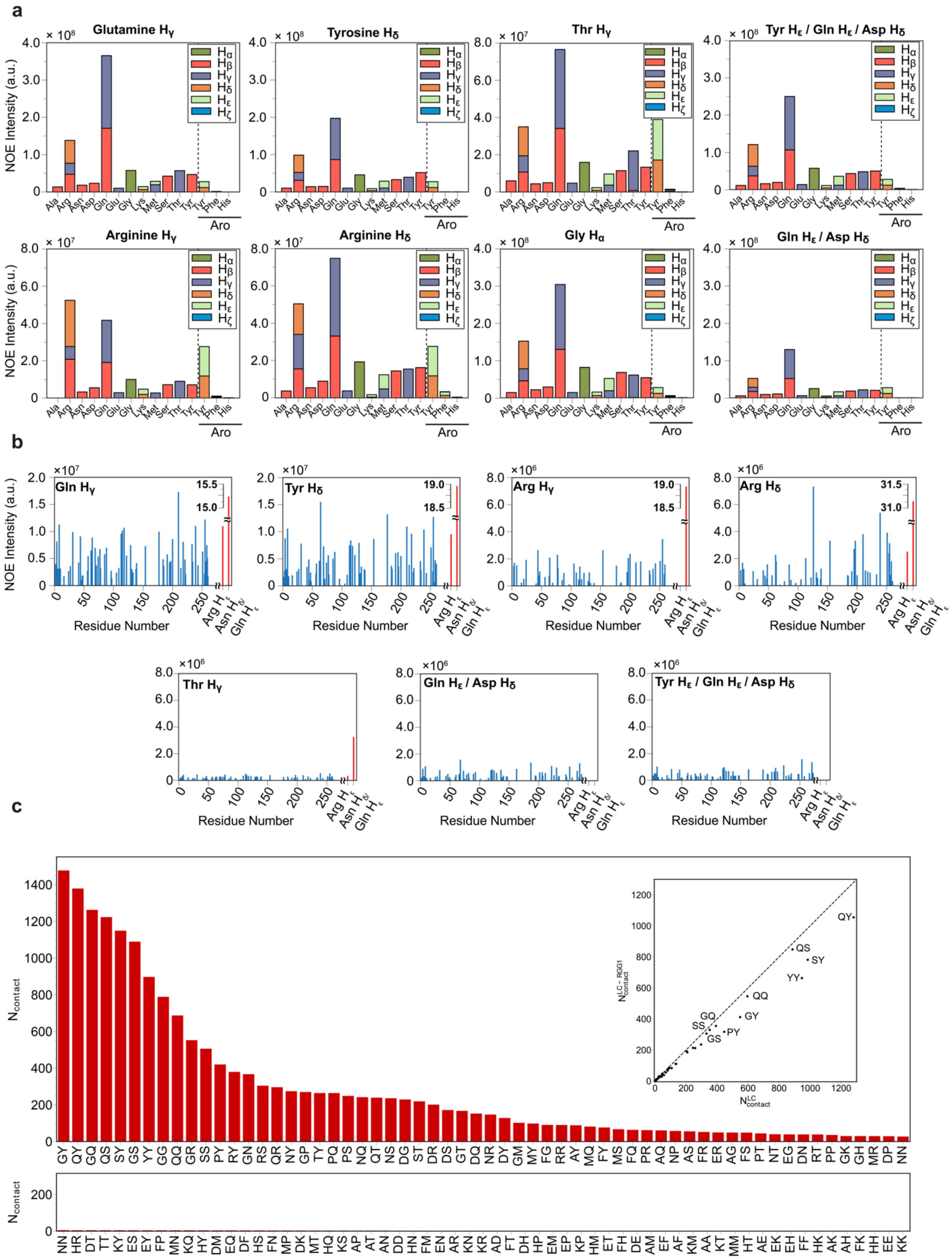


**Figure S2. Intermolecular contacts in the FUS LC-RGG1 condensed phase**

**a** Intermolecular NOE arising from contacts formed in the 1:1 $^{13}C$-$^{15}N$ and $^{12}C^{14}N$ condensed phase samples of FUS LC-RGG1. These data demonstrate that these residue types forms interactions with several other residue types with intensities (partially correlated with residue and pair interaction frequency) which depend on the identity of each residue. Main text figure panels are repeated here to facilitate comparison.
**b** Intermolecular contacts formed from sidechain to backbone positions obtained by analyzing a 1:1 $^{13}C/^{15}N$ and $^{12}C/^{14}N$ condensed phase sample of FUS LC-RGG1. These data demonstrate that sidechain to backbone contacts are distributed across the entirety on the sequence and that arginine $H_\varepsilon$ NOEs are found for some residue types but not others.
**c** Summed and ranked pairwise contacts observed in condensed phase simulations of the LC-RGG1. (inset) Correlation of the pairwise condensed phase contacts formed in the LC condensed phase compared to the LC-RGG1 condensed phase, showing that LC:LC pairs are lower in the LC-RGG1 phase, likely due to LC interaction with RGG1 domains.

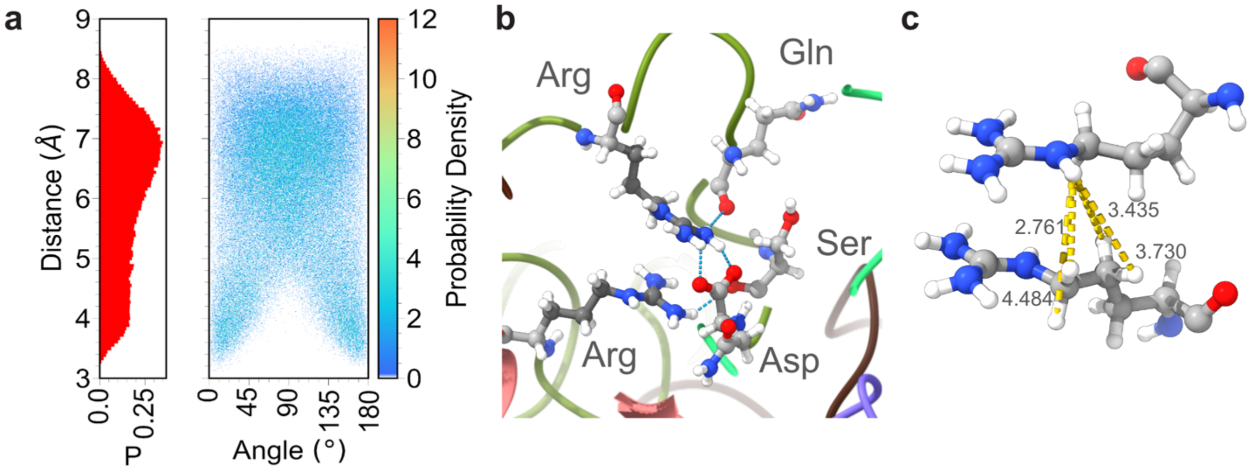


**Figure S3. Arginine-arginine contacts in the FUS LC-RGG1 phase separation**
**a** Simulations of FUS LC-RGG1 show that when arginine's guanidyl groups approach (<6 Å distance between heavy atoms), the center of mass of the guanidyl group can form close interactions with center of mass distance (vertical axis) coming into close proximity (left). Stacking interactions are discernable by the angle distribution of the vector normal to the guanidyl plane, showing bias towards parallel configurations when the guanidyl groups are close.
**b** These arginine-arginine interactions between guanidyl groups can be stabilized by recruitment of negatively charged residues (e.g. Asp) as well as polar residues (e.g. Gln and Ser) that can form salt bridges and hydrogen bonds.
**c** The stacked interactions bring hydrogen positions that show $^{1}H$-$^{1}H$ NOEs in the NMR experiments into close proximity, for example shown here are the short distances (shown in Å) between arginine $H_\varepsilon$ and arginine $H_\delta$ and $H_\gamma$ that can give rise to strong NOEs.

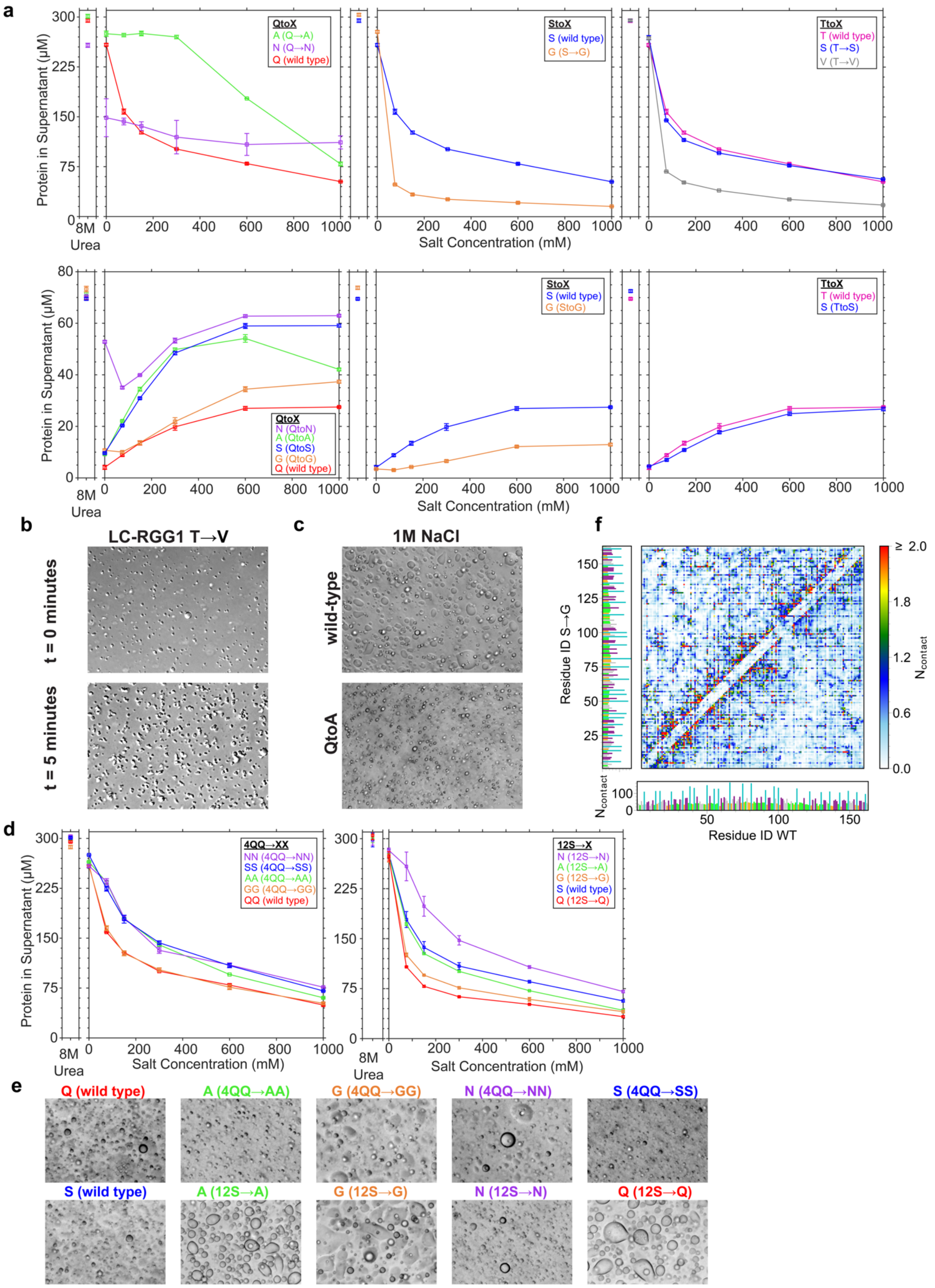
a
Protein in Supernatant (μM)
8M Urea
Salt Concentration (mM)
QtoX
A (Q→A)
N (Q→N)
Q (wild type)
StoX
S (wild type)
G (S→G)
TtoX
T (wild type)
S (T→S)
V (T→V)
QtoX
N (QtoN)
A (QtoA)
S (QtoS)
G (QtoG)
Q (wild type)
StoX
S (wild type)
G (StoG)
TtoX
T (wild type)
S (TtoS)
b
LC-RGG1 T→V
t = 0 minutes
t = 5 minutes
c
1M NaCl
wild-type
QtoA
f
Residue ID S→G
Residue ID WT
$N_{contact}$
≥ 2.0
d
4QQ→XX
NN (4QQ→NN)
SS (4QQ→SS)
AA (4QQ→AA)
GG (4QQ→GG)
QQ (wild type)
12S→X
N (12S→N)
A (12S→A)
G (12S→G)
S (wild type)
Q (12S→Q)
e
Q (wild type)
A (4QQ→AA)
G (4QQ→GG)
N (4QQ→NN)
S (4QQ→SS)
S (wild type)
A (12S→A)
G (12S→G)
N (12S→N)
Q (12S→Q)

**Figure S4. Phase separation of FUS LC and LC-RGG1 is dependent on polar residue composition**

**a** Salt dependent phase diagrams of polar residue substitution variants in the FUS LC and LC-RGG1 sequences. These data are shown as a bar plot for 150 mM NaCl in Figure 4b for variants that show liquid-like droplets. Values for 8M urea are controls to ensure no phase separation (though Q→N still shows some evidence for aggregation or loss by sticking to the tube).

**b** DIC micrographs of T→V in the LC-RGG1 sequence. Spherical droplets are formed at t=0 that quickly proceed to dynamically arrested higher order assemblies after 5 minutes.

**c** DIC micrographs of FUS LC Q→A at 1M NaCl. These images demonstrate that while phase separation is not observed at low salt concentrations at 300 μM FUS LC, formation of liquid phase separated droplets is possible at these protein concentrations at high enough salt concentrations.

**d** Salt dependent phase diagrams of partial polar residue substitution variants in FUS LC. These data are shown as a bar plot for 150 mM NaCl in Figure 4e for variants that show liquid-like droplets. Values for 8M urea are controls to ensure no phase separation (though extinction coefficient for FUS LC-RGG1 in 8M urea is different from in native buffer, so values appear higher than 60 μM concentrations which were measured by dilution in native buffer).

**e** DIC micrographs of the partial polar residue substitution variants. Each sequence formed phase separated droplets.

**f** Interaction profiles comparing the wild type and S→G FUS LC sequences. Interactions across the FUS LC sequence are slightly decreased at the positions changed from serine to glycine but enhanced at adjacent positions.

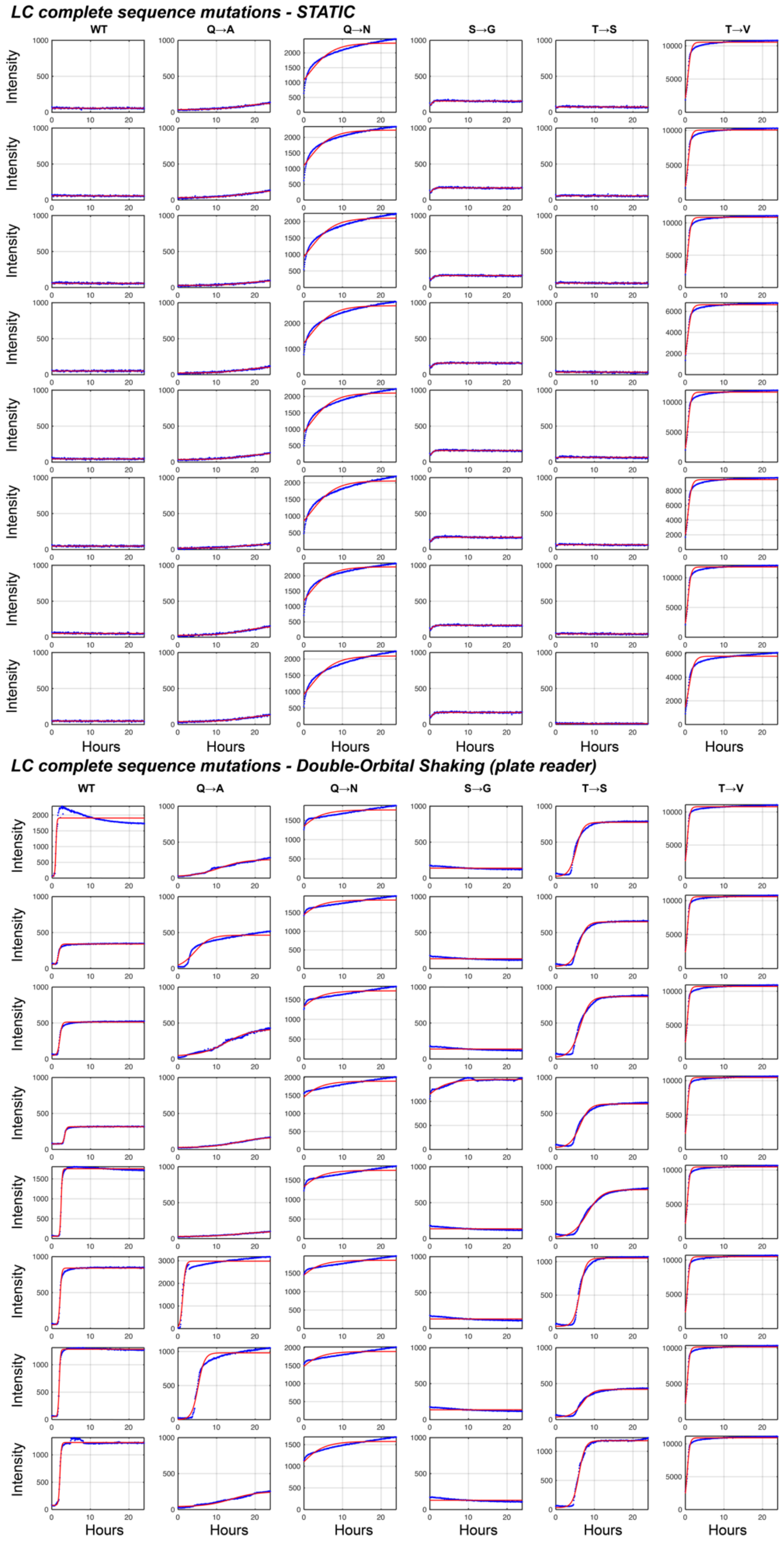


**Figure S5.** Individual replicates and sigmoid fits of the ThT fluorescence assay probing formation of ThT-positive aggregation of the wild type LC domain and complete replacement substitution variants under quiescent and double orbital shaking conditions.

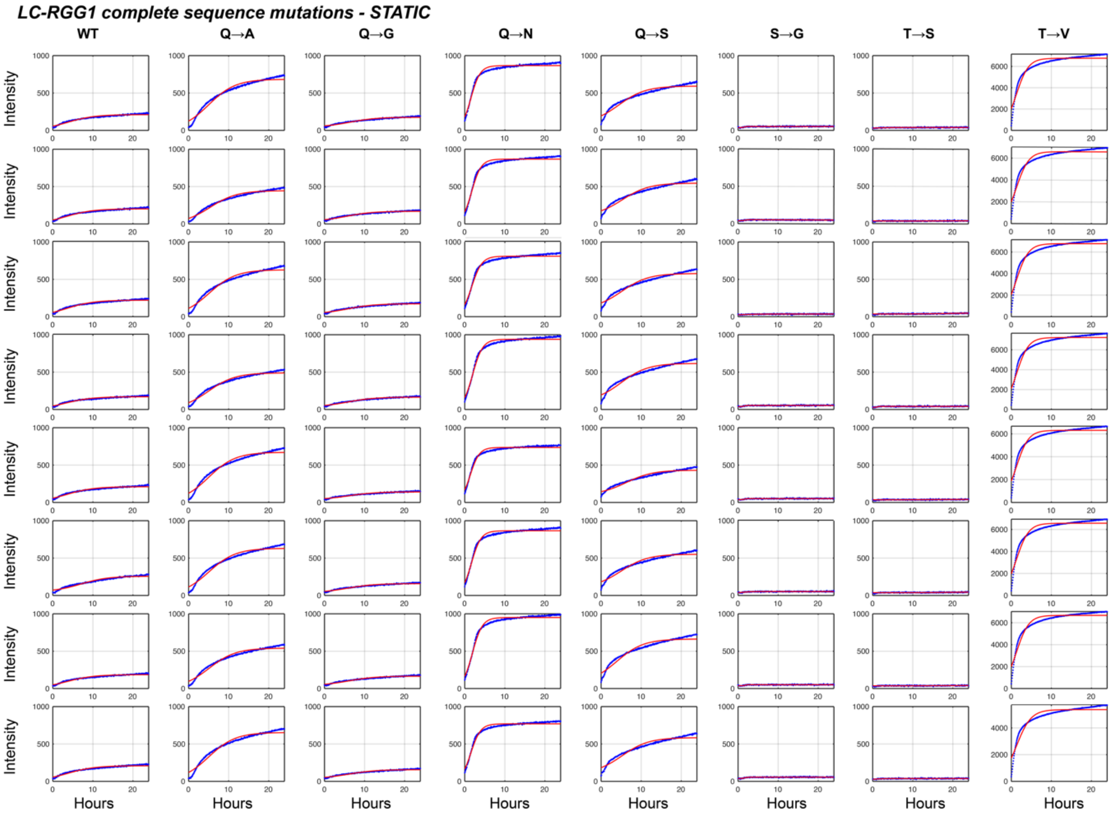


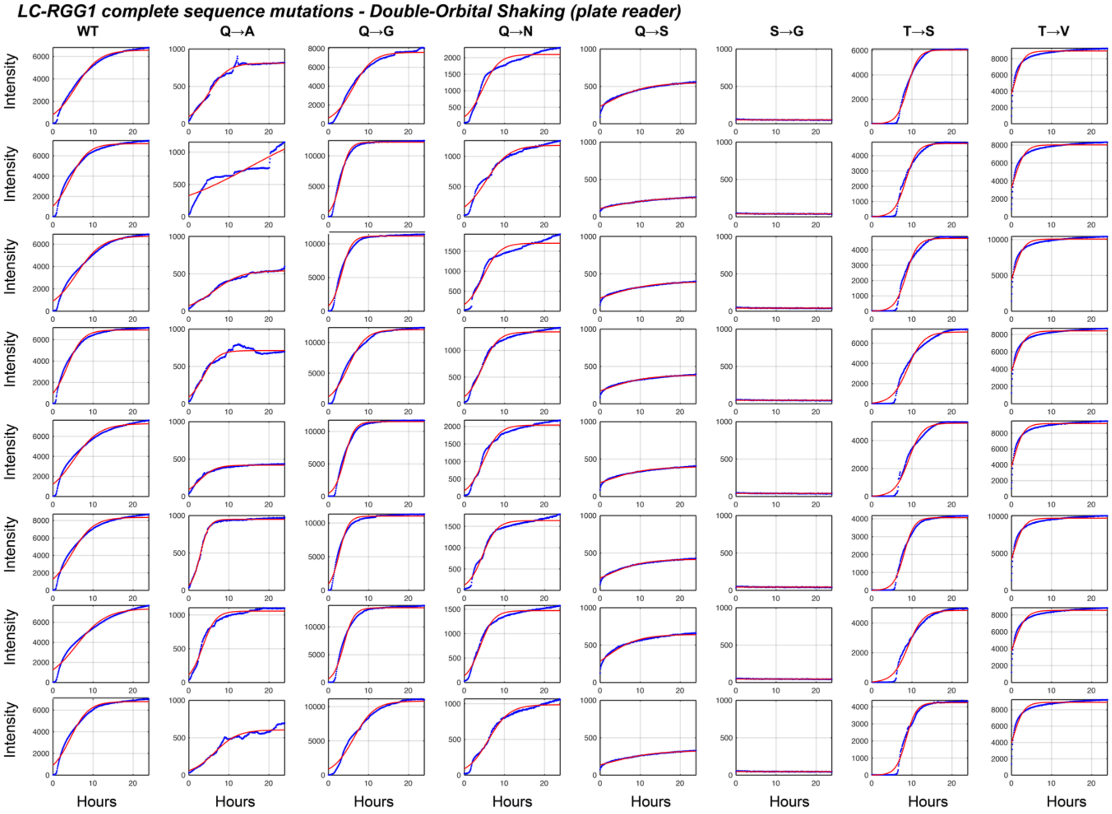


**Figure S6.** Individual replicates and sigmoid fits of the ThT fluorescence assay probing formation of ThT-positive aggregation of the wild type LC-RGG1 domain and complete replacement substitution variants under quiescent and double orbital shaking conditions.

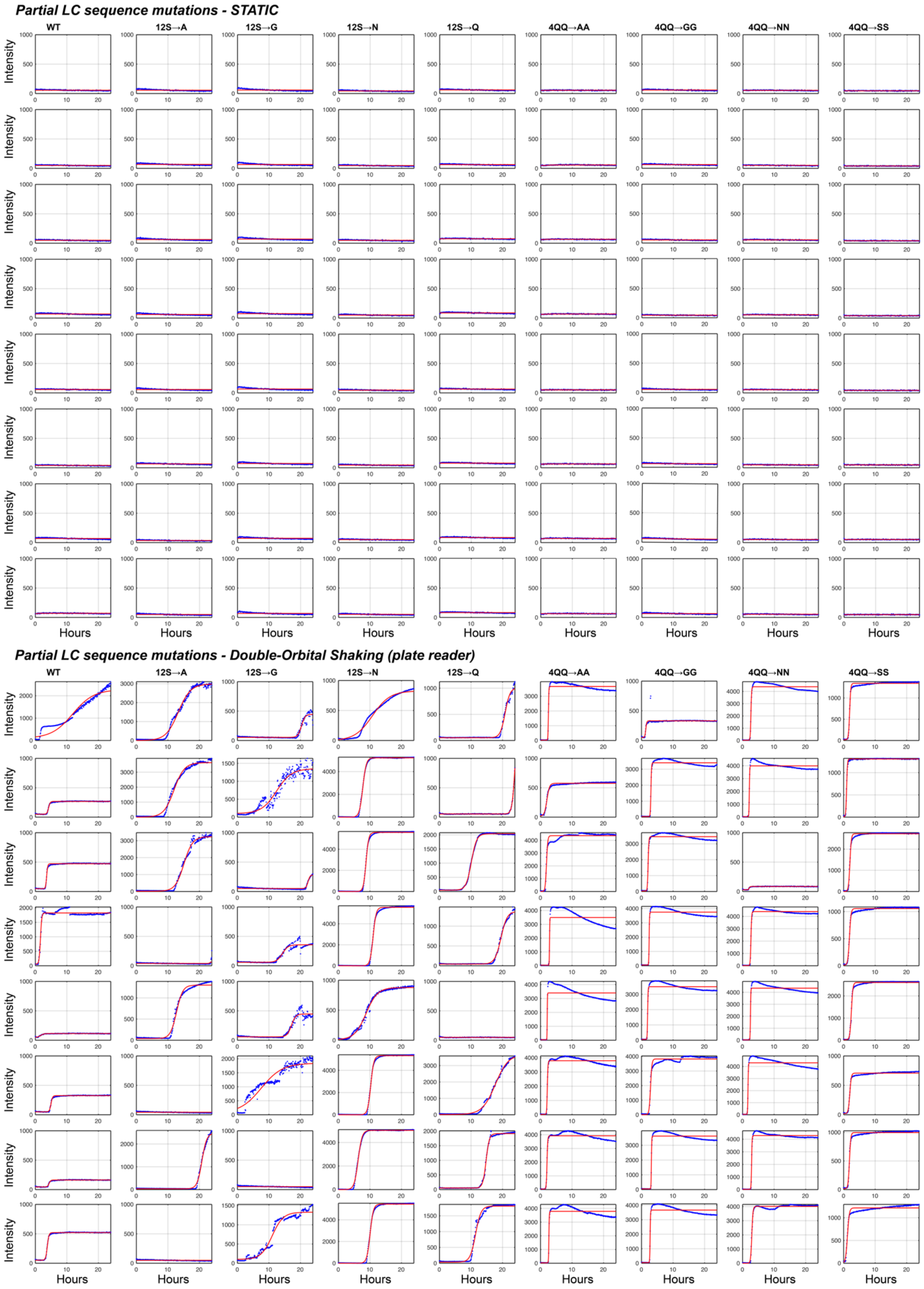


**Figure S7.** Individual replicates and sigmoid fits of the ThT fluorescence assay probing formation of ThT-positive aggregation of the wild type LC domain and partial substation variants (12S→X and 4QQ→XX) under quiescent and double orbital shaking conditions.

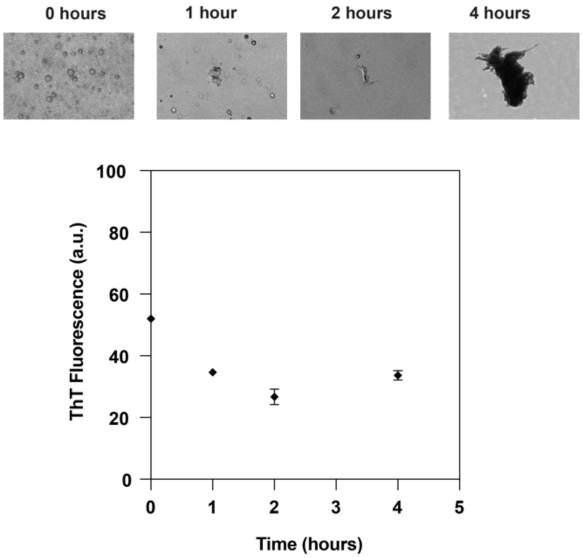


**Figure S8. Not all irregular shaped aggregates show ThT fluorescence enhancement despite liquid-to-solid type transition**
(Top) Brightfield microscopy of FUS LC 4QQ→NN samples under quiescent conditions with 20 μM Thioflavin T over the course of 4 hours. (Bottom) ThT fluorescence for these samples over the 4 hour time course. ThT fluorescence enhancement is not observed for these samples despite clear conformational change from liquid-like droplets to irregularly shaped aggregates.

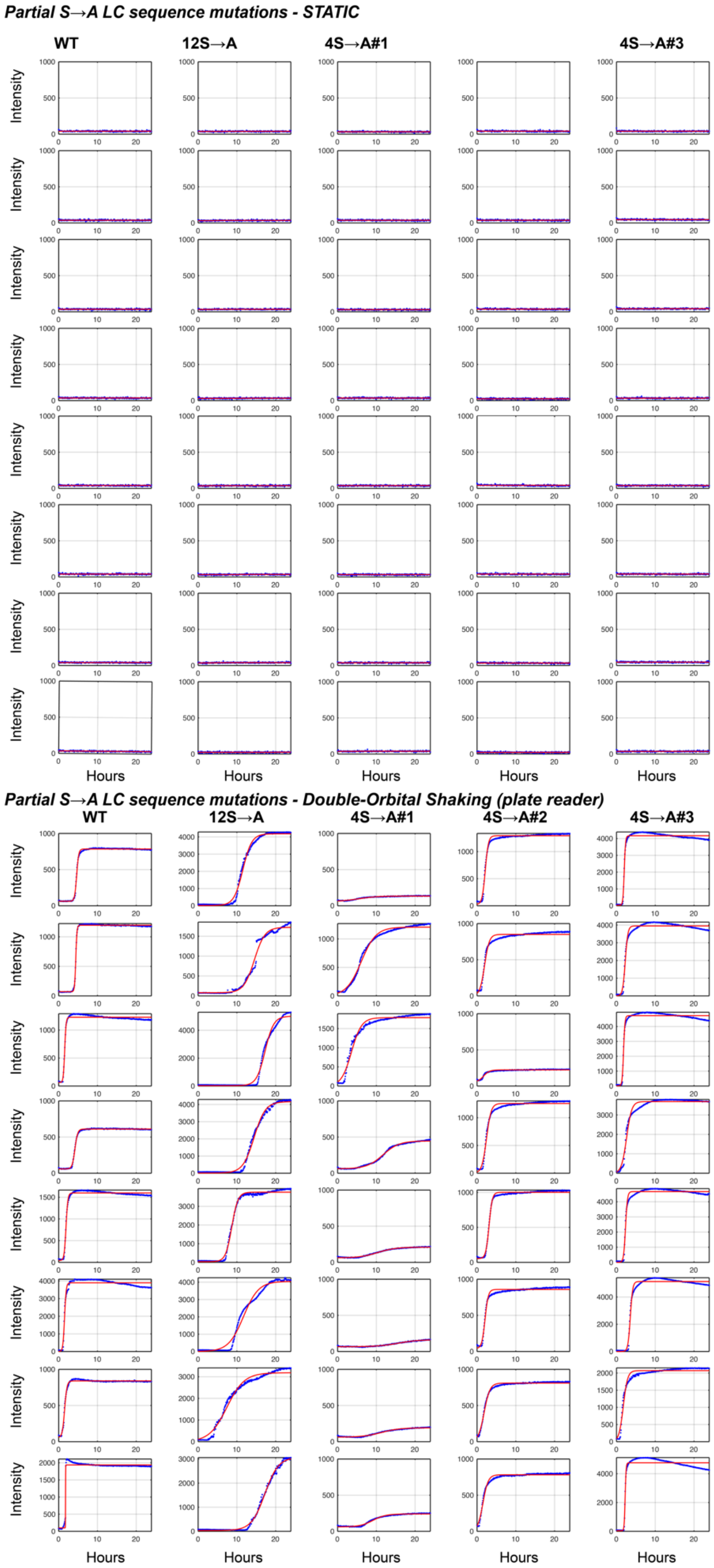


**Figure S9.** Individual replicates and sigmoid fits of the ThT fluorescence assay probing formation of ThT-positive aggregation of the wild type LC domain and the serine to alanine variant series under quiescent and double orbital shaking conditions.
